# Comb-structured mRNA vaccine tethered with short double-stranded RNA adjuvants maximizes cellular immunity for cancer treatment

**DOI:** 10.1101/2022.01.18.476829

**Authors:** Theofilus A. Tockary, Saed Abbasi, Miki Matsui-Masai, Naoto Yoshinaga, Eger Boonstra, Zheng Wang, Shigeto Fukushima, Kazunori Kataoka, Satoshi Uchida

## Abstract

Integrating antigen-encoding mRNA and immunostimulatory adjuvant into a single formulation is a promising approach to potentiating the efficacy of mRNA vaccines. Here, we developed a scheme based on RNA engineering to integrate adjuvancy directly into antigen-encoding mRNA strands without hampering the ability to express antigen proteins. Short double-stranded RNA (dsRNA) was designed to target retinoic acid-inducible gene-I (RIG-I), an innate immune receptor, for effective cancer vaccination and then tethered onto mRNA strand via hybridization. Tuning the dsRNA structure and microenvironment by changing its length and sequence enabled the determination of the structure of dsRNA-tethered mRNA efficiently stimulating RIG-I. Eventually, the formulation loaded with dsRNA-tethered mRNA of the optimal structure effectively activated mouse and human dendritic cells and drove them to secrete a broad spectrum of proinflammatory cytokines without increasing the secretion of anti-inflammatory cytokines. Notably, the immunostimulating intensity was tunable by modulating the number of dsRNA along mRNA strand, which prevents excessive immunostimulation. Versatility in the applicable formulation is a practical advantage of the dsRNA-tethered mRNA. Its formulation with three existing systems, *i*.*e*., anionic lipoplex, ionizable lipid-based lipid nanoparticles, and polyplex micelles, induced appreciable cellular immunity in the mice model. Of particular interest, dsRNA-tethered mRNA encoding ovalbumin (OVA) formulated in anionic lipoplex used in clinical trials exerted a significant therapeutic effect in the mouse lymphoma (E.G7-OVA) model. In conclusion, the system developed here provides a simple and robust platform to supply the desired intensity of immunostimulation in various formulations of mRNA cancer vaccines.

## Introduction

Messenger RNA (mRNA) vaccines have promising potential in cancer immunotherapy, with numerous clinical trials in progress (1-3). Advantages of mRNA in cancer vaccination include efficient induction of cytotoxic T-lymphocyte (CTL) immune responses via major histocompatibility complex (MHC) class I pathway and flexible designing for targeting neo-antigens just by changing mRNA sequences (4, 5). Consequently, various formulations for mRNA vaccine delivery have been developed to maximize antigen expression efficiency. In parallel, vigorous efforts have been devoted to designing safe and effective immunostimulatory adjuvants for robust immunization in cancer vaccines over the past years (2, 6, 7).

Numerous studies have demonstrated the benefit of co-loading antigen and immunostimulatory adjuvant into a single formulation, which ensures co-delivery of these two essential components to the same antigen-presenting cells (8-10). The orthodox approach is to co-load adjuvant molecules with antigen mRNA into the same formulation using proper packaging materials (11, 12). An alternative approach that has emerged recently is to directly integrate adjuvancy into the packaging materials used for formulating mRNA (13-15). Ionizable lipids used for formulating lipid nanoparticles (LNPs) are typical examples of such materials with intrinsic immunostimulatory properties (16-18). Adjuvancy integration into packaging materials instead of using separate adjuvant molecules is beneficial for formulating clinically-translatable vaccines because of the simplicity in the production process (19). Nevertheless, balancing the delivery efficiency and adjuvant activity is still challenging in this strategy, requiring elaborative optimization processes to prepare the packaging materials with desired functionalities. This issue motivated us to develop a novel and simple methodology to maximize the adjuvancy of mRNA formulation just by using existing packaging materials already optimized in terms of delivery efficiency.

A key concept in this newly-developed methodology is incorporating adjuvancy into mRNA itself rather than packaging materials, thereby maximizing immunostimulating and delivery efficiencies of mRNA formulations. For this purpose, comb-structured mRNA was designed by hybridizing antigen-encoding mRNA with immunostimulatory double-stranded RNA (dsRNA) teeth (**Figure 1**). The dsRNA teeth activate innate immune receptors, functioning as adjuvants. This comb-structured mRNA can be encapsulated into various existing mRNA vaccine carriers without changing their original properties and functionalities. Furthermore, changing tooth number enables the immunostimulation intensity of comb-structured mRNA to the optimal extent for each formulation prepared from different packaging materials. Notably, this methodology only requires adding a small amount of dsRNA, a nature-derived safe material, for potentiating existing mRNA vaccines. This feature allows for smooth clinical translation from a safety viewpoint.

**Figure 1.**
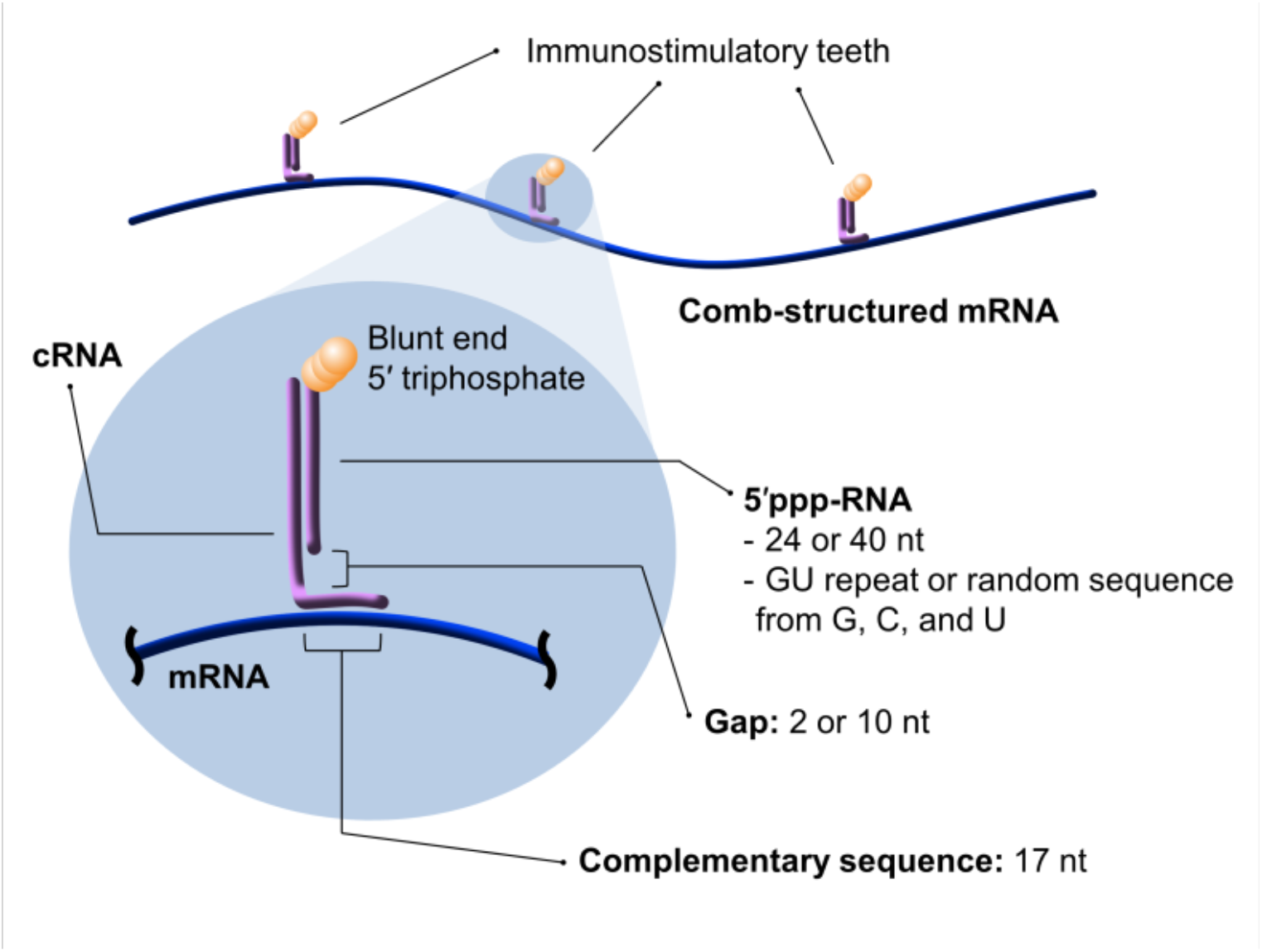
Comb-structured mRNA with immunostimulatory dsRNA teeth. This formulation is prepared from *in vitro* transcribed RNA with 5’ triphosphate (5’ppp-RNA), mRNA, and a chemically synthesized counter RNA strand complementary to the 5’ppp-RNA and particular sequences in mRNA (cRNA). dsRNA teeth from 5’ppp-RNA and cRNA possess blunt-ended 5’ triphosphate, a substrate of RIG-I. Length of cRNA sequence complementary to mRNA is fixed to 17 nt to maintain mRNA translational activity. In addition, design parameters of comb-structured mRNA, including the lengths and sequences of dsRNA teeth, and lengths of gap sequence in cRNA between two regions complementary to 5’ppp-RNA and mRNA, were optimized.

Toward effective cancer immunotherapy, dsRNA teeth were designed to activate retinoic acid-inducible gene-I (RIG-I), an innate immune receptor inducing strong CTL responses (20, 21). RIG-I requires only short dsRNA with 20 base pairs (bp) for activation (22), minimally influencing the total RNA dose. Notably, this dsRNA-toothed approach has the versatility in fine-tuning the dsRNA structure and microenvironment by simply changing its length and sequence and a gap sequence length between dsRNA and mRNA (**Figure 1**). A thorough examination of these variants specified the optimized comb-structure formulation for efficient and specific RIG-I stimulation. Ultimately, the comb-structured mRNA potentiated the following three existing mRNA vaccine systems, possessing completely different immunogenic properties, to successfully induce CTL immune responses: Anionic lipoplex comparable with that used in clinical trials (23, 24), ionizable lipid-based LNP (iLNP), a prevalent platform of current mRNA vaccines (25), and polyplex micelle, a representative polymer-based delivery systems (26).

## Results

### In vitro screening identifies highly immunostimulating comb-structured mRNA

RIG-I prefers dsRNA with 5’ triphosphate (5’ppp) at the blunt end as a substrate (22, 27). For the preparation of comb-structured mRNA stimulating RIG-I, we designed a short RNA with 5’ triphosphate (5’ppp-RNA) and a counter RNA strand complementary both to this 5’ppp-RNA and the particular sequences in mRNA (cRNA) (**Figure 1**). While *in vitro* transcription (IVT) without a 5’ cap analog is a cost-effective method to prepare 5’ppp-RNA, there is an issue of producing contaminant RNA strand complementary to the intended RNA strand mainly via RNA-templated RNA transcription (28-30). Contaminant dsRNA formed from this complementary RNA may negatively influence the hybridization process and immunostimulatory property of the RNA. Here, we circumvented this issue by designing the sequence of 5’ppp-RNA to contain no adenine (A) residue and several uracil (U) residues. IVT of 5’ppp-RNA without A residue circumvents the formation of complementary contaminant RNA, which should have several A residues in its sequence. The risk of complementary RNA formation may be further reduced by avoiding self-complementary sequences in 5’ppp-RNA, as RNA hybridization to self-RNA in cis or trans through self-complementary sequences plays a significant role in complementary RNA synthesis (31). Accordingly, as for 5’ppp-RNA, we prepared 24 nucleotides (nt) or 40 nt GU-repeat RNA, lacking a self-complementary sequence. We also prepared RNA with a random sequence from guanine (G), cytosine (C), and U residues containing self-complementary sequences as control (**Supplementary Table S1**). cRNA was chemically synthesized to avoid the formation of IVT by-products (**Supplementary Table S2**). The length of the complementary sequence in cRNA hybridized to mRNA was fixed to 17 nt because both translational activity and immunogenicity of mRNA were still preserved even after hybridization of 17 nt complementary RNA (32). We placed 2 nt or 10 nt gap sequence between complementary sequences to 5’ppp-RNA and mRNA in the cRNA strand, hypothesizing that such a difference in gap sequence length might influence the microenvironment of dsRNA teeth for recognition by innate immune receptors. *Gaussia luciferase* (*gLuc*) mRNA was used as a standard to quantify the protein translational activity of comb-structured mRNA. Successful hybridization was confirmed by ultrafiltration, which separates free-formed dsRNA passing through the filter from those hybridized with mRNA using Cy5-labeled dsRNA. Eventually, in this experiment, Cy5-fluorescence was undetected in the flow-through, showing that almost all Cy5-labeled dsRNA was successfully hybridized to mRNA (**Supplementary Figure S1**).

There are several design parameters in immunostimulatory dsRNA teeth, including the lengths and sequences of dsRNA and the lengths of gap sequence in cRNA between two regions complementary to 5’ppp-RNA and mRNA (**Figure 1**). These parameters were optimized based on the immunostimulating property of comb-structured mRNA after its introduction to DC2.4 cells, a murine DC-derived cell line. Lipofectamine LTX, a commonly used lipid-based transfection reagent, was used for mRNA introduction, as its efficient mRNA introduction capability *in vitro* allows for high-throughput evaluation of immunostimulatory and translational properties of mRNA. Notably, the formulation materials used in Lipofectamine LTX exert negligible immunostimulatory properties. Actually, as shown in Figures 2 and 5, Lipofectamine LTX loading mRNA without immunostimulatory dsRNA tooth showed a negligible immunostimulating effect *in vitro*. Thus, Lipofectamine LTX formulation is suitable for examining the inherent immunostimulatory and translational properties of loaded mRNA with varying structures of dsRNA teeth without bias. Transcripts of proinflammatory genes, *i*.*e*., *interferon β* (*IFN-β*) and *interleukin 6* (*IL-6*), were quantified 4 h after transfection to evaluate the immunostimulatory property of mRNA with dsRNA teeth.

**Figure 2.**
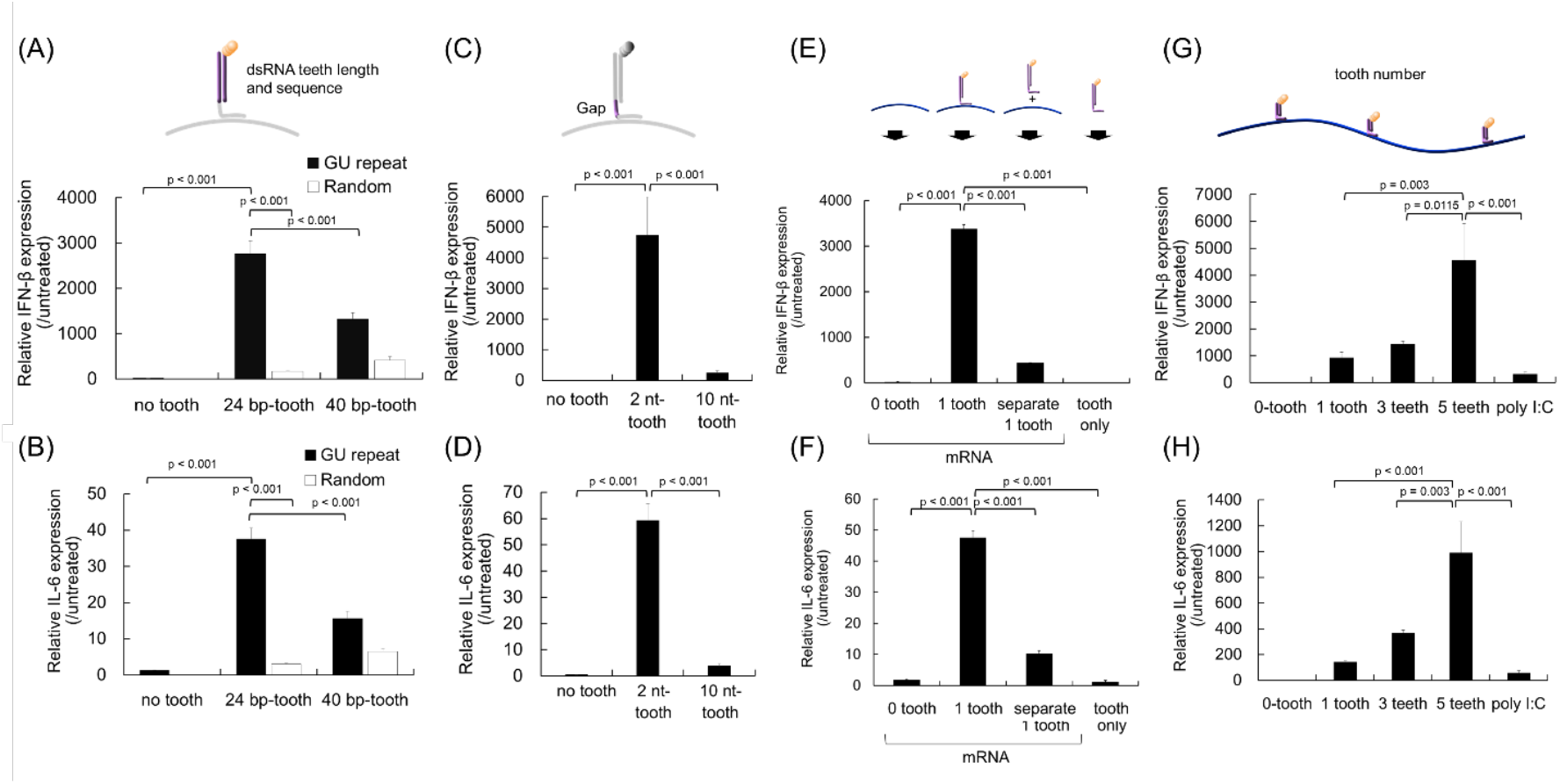
Optimization of tooth design for efficient immunostimulation. Transcript levels of *interferon (IFN)-β* (**A, C, E, G**) and *interleukin (IL)-6* (**B, D, F, H**) were measured using quantitative PCR 4 h after mRNA introduction to DC2.4 cells. (**A, B**) Effect of dsRNA lengths and sequences. (**C, D**) Effect of gap RNA lengths in cRNA between two regions complementary to 5’ppp-RNA and mRNA. (**E, F**) Effect of dsRNA tooth and mRNA co-delivery method. Introduction of 24 nt GU-repeat tooth alone and separate introduction of mRNA and 24 nt GU-repeat tooth were performed. **(G, H**) Effect of tooth numbers. The teeth used possessed dsRNA with 24 bp GU-repeat and a 2 nt gap (dsRNA24-GUrepeat/2-gap). *n* = 5 in (**A-D**), *n* = 6 in (**E-H**).

Firstly, we optimized sequences and lengths in the dsRNA tooth using comb-structured mRNA with one tooth. GU-repeat tooth induced enhanced expression of proinflammatory transcripts compared to random sequence (**Figure 2A,B**), possibly because GU-repeat sequence might avoid the formation of complementary RNA strand by-products as aforementioned. Elongation of GU-repeat tooth from 24 bp to 40 bp resulted in a slight reduction in proinflammatory transcripts. Although the mechanism underlying this result is unclear, longer complementary RNA strands might have more freedom to hybridize with each other at unintended positions, hampering the formation of intended dsRNA possessing 5’-triphosphate at the blunt end, a preferred ligand of RIG-I. Then, gap RNA length in cRNA between two regions complementary to 5’ppp-RNA and mRNA was optimized for the dsRNA tooth with 24 bp GU-repeat. As seen in **Figure 2C,D**, the 2 nt gap revealed higher proinflammatory transcripts levels than the 10 nt gap. Presumably, gap length might alter the local environment of dsRNA and, as a result, affect the process of dsRNA recognition by innate immune receptors. Interestingly, the introduction of only a free-formed 24 bp GU-repeat tooth and the separate introduction of mRNA and 24 bp GU-repeat tooth failed to induce strong immunostimulation (**Figure 2E,F**). The microenvironment of dsRNA in these groups might also be different from that of dsRNA tethered to mRNA strands with a 2 nt gap in cRNA. This result suggests the significance of hybridizing dsRNA to mRNA strands for stimulating innate immunity. Accordingly, we selected to use dsRNA with 24 bp GU-repeat and a 2 nt gap (dsRNA24-GUrepeat/2-gap) in the following experiments. As seen in **Figure 2G,H**, expression levels of proinflammatory transcripts increased with the number of dsRNA24-GUrepeat/2-gap teeth from 1 to 5.

### Comb-structured mRNA activates RIG-I for immunostimulation

Several innate immune receptors potentially recognize comb-structured mRNA. They include dsRNA receptors (RIG-I (22), melanoma differentiation-associated gene (MDA)-5 (33), Toll-like receptor (TLR) 3 (34)), and a single strand RNA receptor (TLR7 (35)). RIG-I involvement in inflammatory response due to mRNA transfection was evaluated using the *RIG-I* knockout RAW-Lucia cell. RAW-Lucia cell is a macrophage-derived cell line genetically modified to express Lucia luciferase (lLuc) responding to proinflammatory stimuli under the promoter responsive to interferon regulatory factors. As seen in **Figure 3A**, without *RIG-I* knockout (WT), comb-structured mRNA with 1, 3, and 5 teeth showed enhanced lLuc expression in RAW-Lucia cells compared to untreated control and mRNA without a tooth. In contrast, comb-structured mRNA with 1, 3, and 5 teeth did not increase lLuc expression levels in *RIG-I* knockout cells. Further notably, without 5’ppp, a critical motif in dsRNA for RIG-I recognition (22), lLuc expression levels were comparable between comb-structured mRNA and untreated control in RAW-Lucia cells without *RIG-I* knockout. These results highlight a pivotal role of RIG-I in the immunostimulation induced by comb-structured mRNA. On the contrary, knockout of *MDA-5* in RAW Lucia cells showed almost no influence on lLuc expression after treatment with comb-structured mRNA. This result suggests that comb-structured mRNA possessing dsRNA24-GUrepeat/2-gap teeth does not stimulate MDA-5, which requires dsRNA longer than 2 kb for its stimulation (33).

**Figure 3.**
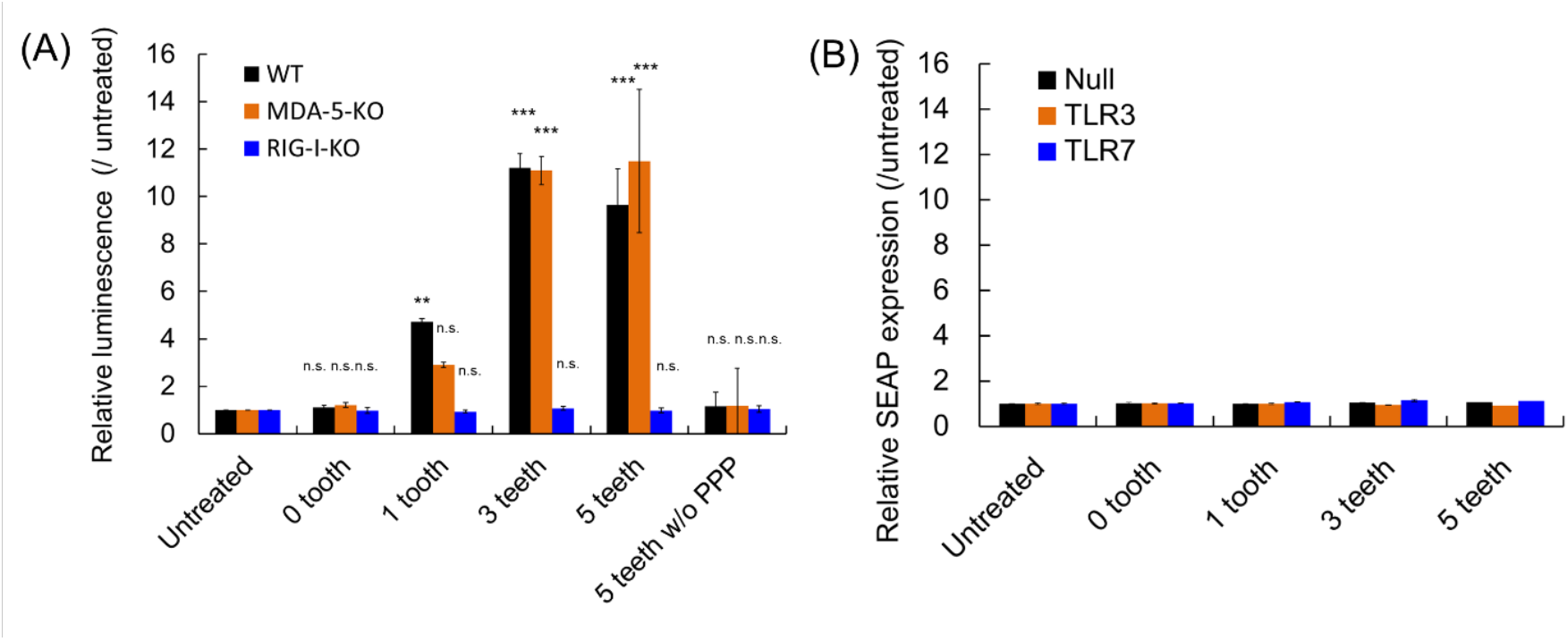
Contribution of innate immune receptors for immunostimulation by mRNA. (**A**) Comb-structured mRNA was added to RAW cells without knockout of immune receptors (WT) and with *MDA-5* or *RIG-I* knockout (MDA-5-KO, RIG-I-KO). The cells were genetically modified to express Lucia luciferase (lLuc) after proinflammatory stimulation for quantifying immunostimulation intensity based on lLuc expression. *n* = 6. (**B**) Comb-structured mRNA was added to HEK 293 cells, genetically modified to express TLR3 or TLR7, or without transformation to express TLRs (Null). The cells were transformed to express secreted embryonic alkaline phosphatase (SEAP) reporter after nuclear factor-κB (NF-κB) stimulation. *n* = 6. Abbr.: PPP, 5’ triphosphate. **, p < 0.01; ***, p < 0.001, n.s., non-significant versus untreated, respectively.

The involvement of TLRs was studied using a human embryonic kidney-derived cell line, HEK293 cells, genetically modified to express human TLR3 or TLR7, denoted as HEK-TLR3 or HEK-TLR7, respectively. Note that original HEK 293 cells, denoted as HEK-null, exhibited a negligible level of TLR expression. These three cell lines were further transformed to express secreted embryonic alkaline phosphatase (SEAP) reporter after nuclear factor-κB (NF-κB) stimulation. In HEK-TLR3 and HEK-TLR7, mRNA with 1, 3, and 5 teeth and without teeth provided SEAP expression levels comparable with those in untreated control, indicating a minimal role of TLR3 and 7 in immunostimulation by comb-structured mRNA (**Figures 3B**).

### Comb-structured mRNA efficiently activates DCs with minimal influence on translational activity

Detailed functional analyses of comb-structured mRNA were performed using mouse primary bone marrow-derived dendritic cells (BMDCs). First, the capability of comb-structured mRNA for activating BMDCs was evaluated by measuring expression levels of surface marker proteins, including CD86, CD40, and MHC class I and II (MHC I and MHC II). After introducing into BMDCs, mRNA with a tooth of dsRNA24-GUrepeat/2-gap significantly increased CD86, CD40, and MHC I expression levels compared to mRNA without tooth (**Figures 4A-G**), demonstrating efficient activation of BMDCs by comb-structured mRNA. In most cases, these markers’ expression level tends to become maximal at the three teeth. Meanwhile, the teeth introduction into mRNA has minimal effect on MHC II expression (**Figure 4H**). Notably, compared to polyinosinic-polycytidylic acid (poly I:C), mRNA possessing 1 – 5 teeth showed enhanced CD86, CD40, and MHC I expression in BMDCs, demonstrating intense activity of comb structure as an immunostimulatory adjuvant.

**Figure 4.**
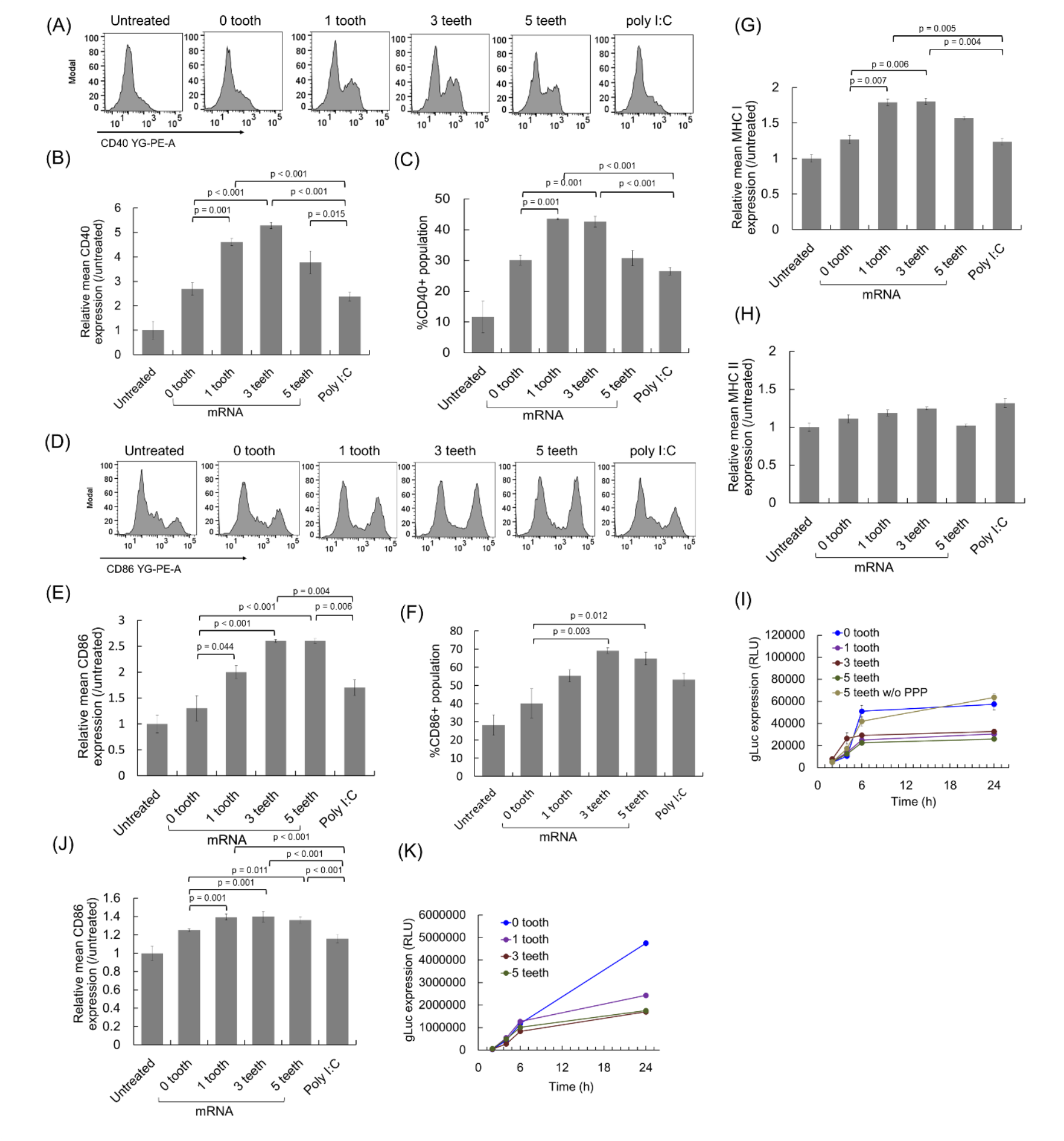
Activation of dendritic cells by comb-structured mRNA. mRNA was added to mouse BMDCs (**A-I**) and human BMDCs (**J, K**). (**A-H, J**) Expression of surface markers, CD40 (**A-C**), CD86 (**D-F, J**), MHC I (**G**), and MHC II (**H**) was quantified using immunocytochemistry 24 h after mRNA addition. *n* = 4 (**I, J**) Expression of gLuc was measured using the cultured medium. *n* = 6. Abbr.: PPP, 5’ triphosphate.

Then, the efficiency of antigen expression was studied using gLuc as a reporter. gLuc expression efficiency of comb-structured mRNA with 1, 3, and 5 teeth was preserved to approximately 50% of the efficiency of mRNA without a tooth in BMDCs (**Figure 4I**). According to previous studies, activation of innate immune responses harms translational activity of mRNA (36, 37), which may reduce gLuc expression efficiency after teeth attachment in the present experiment. To study this issue, we introduced *gLuc* mRNA with 5 teeth without 5’ppp, which lacks the immunostimulating ability (**Figure 3**). Notably, mRNA having 5 teeth without 5’ triphosphate exerted similar gLuc expression to mRNA without a tooth (**Figure 4I**). These results indicate that the reduction of translational activity after the teeth attachment is attributed to the induction of innate immune responses rather than the presence of teeth on the mRNA strand. This finding is consistent with our previous result that mRNA preserves translational activity after hybridization with 17 nt complementary RNA (32).

The feasibility of comb-structured mRNA was also studied for future clinical translation using human DCs. The introduction of 1, 3, and 5 teeth significantly improved the CD86 expression levels compared to mRNA without tooth and poly I:C (**Figure 4J**). Furthermore, in a reporter assay using *gLuc* mRNA, comb-structured mRNA preserved approximately half of the translational activity of mRNA without tooth (**Figure 4K**). These results indicate the availability of comb-structured mRNA in human antigen-presenting cells.

### Comb-structured mRNA broadly stimulates the expression of proinflammatory molecules

Further immunological characterization of comb-structured mRNA was performed using multiplex immunoassay and enzyme-linked immunosorbent assay (ELISA) to measure protein expression levels of 25 types of cytokines, interferons, and chemokines. Twenty-four hours after mRNA introduction to mouse BMDCs, the concentration of these immune molecules in the culture medium was measured (**Figure 5, Supplementary Figure S2**). The fold-change compared to untreated control is shown as a color gradient in **Figure 5** for 16 of the 25 immune molecules, while expression levels of the remaining nine molecules were below the detection limit.

**Figure 5.**
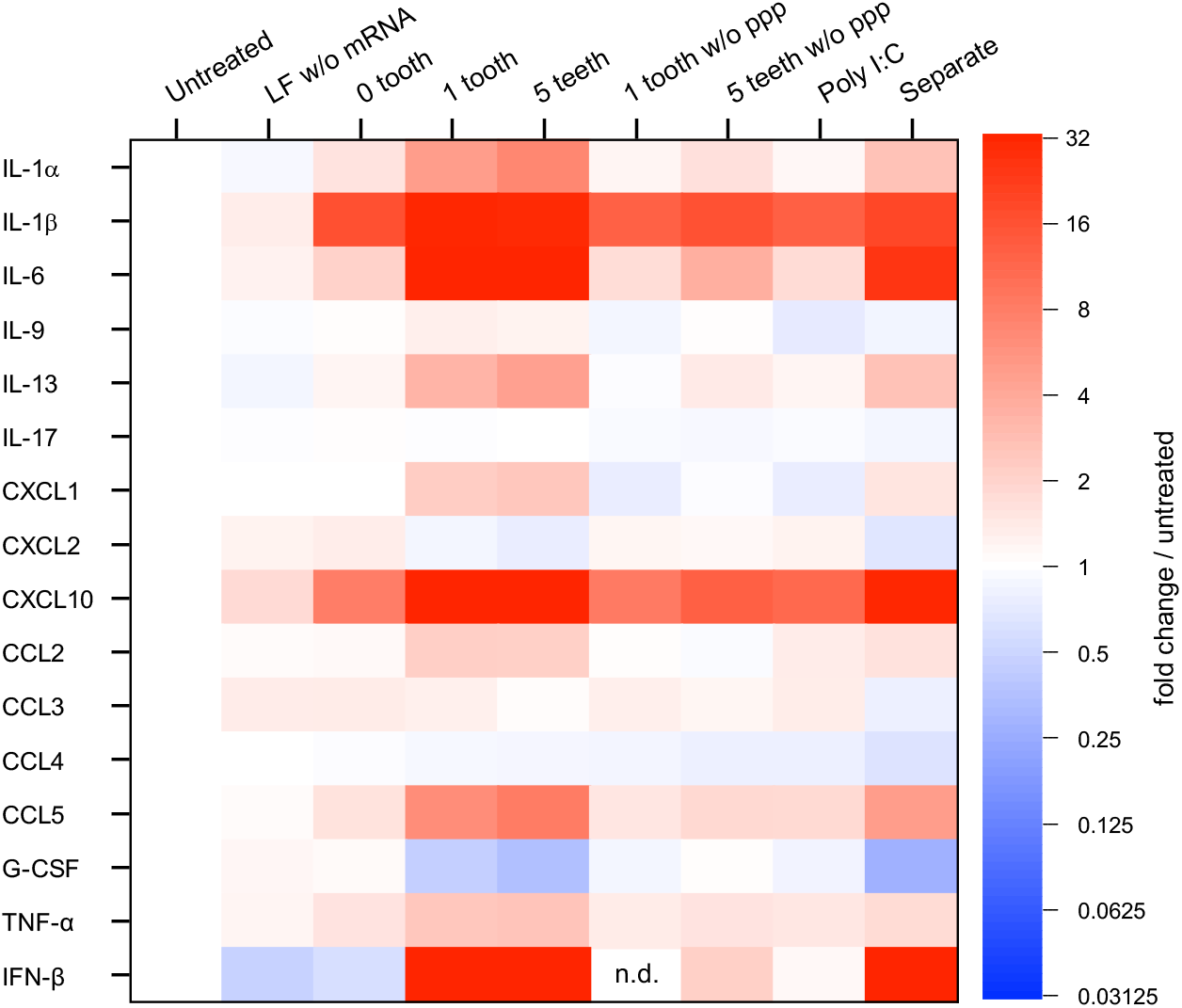
Immunological profiling of BMDCs after mRNA treatment. Protein expression levels of 25 types of cytokines, interferons, and chemokines were quantified 24 h after mRNA introduction to mouse BMDCs. Colors in the heatmap represent protein levels relative to untreated control. Among 25 types of molecules, nine types (IFN-γ, IL-2, 4, 5, 7, 10, 12p40, 12p70, 15) were below detection limits, and data from the other 16 are shown. *n* = 6. Abbr.: LF, lipofectamine; PPP, 5’ triphosphate; G-CSF, granulocyte colony-stimulating factor. In *Separate*, mRNA and tooth with tooth amount equal to that in *1 tooth* were separately formulated with LF for addition to BMDCs.

Two proinflammatory cytokines (interleukin (IL)-1β, IL-6) and one chemokine (C-X-C motif chemokine ligand (CXCL)10) expressed more than 2-fold for mRNA without tooth compared to the untreated control and the control treated with lipofectamine without mRNA (**Figure 5**). This result revealed the immunostimulatory property of mRNA. Intriguingly, compared to mRNA without a tooth, comb-structured mRNA further increased the expression of IL-1β by approximately 2-fold, IL-6 by 17-33-fold, and CXCL10 by 5-7-fold, in a manner dependent on the tooth number. Further notably, secretion of several proinflammatory molecules was enhanced only by comb-structured mRNA, not by mRNA without a tooth. Comb-structured mRNA enhanced the expression levels an approximately 100-fold for a type I interferon (IFN-β), and 2-10-fold for two proinflammatory cytokines (IL-1α, tumor necrosis factor (TNF)-α) and three chemokines (CXCL1, CC chemokine ligand (CCL)2, CCL5) compared to untreated control. In contrast, mRNA without teeth did not increase the expression of these immune molecules by more than 2-fold. These results show that the teeth tethering to the mRNA strand reinforces the immunostimulatory property of mRNA and broadens the types of secreted immune molecules. Such reinforcement was not observed for the mRNA tethering the teeth lacking a 5’ppp group, indicating the involvement of RIG-I in this immunostimulation process. The separate introduction of mRNA and dsRNA also appears to induce relatively high levels of proinflammatory proteins in the heatmap. However, the levels were lower than observed after the introduction of comb-structured mRNA, even when the amount of dsRNA was equal between these two groups (**Supplementary Figure S2**). This result is consistent with the quantification of IFN-β and IL-6 at the mRNA level (**Figure 2E,F**). Despite the properties of comb-structured mRNA to broadly stimulate the secretion of proinflammatory cytokines, *i*.*e*., type I interferon and chemokines, it provided a modest influence on the expression of anti-inflammatory cytokines (IL-10) and T helper 2 (Th2)-related cytokines (IL-4, IL-5, IL-9, IL-13) (38). The expression was undetected for IL-4, IL-5, and IL-10, low for IL-13, and unchanged for IL-9 after the comb-structure installation. These immunological profilings also indicate that comb-structured mRNA exerts much stronger immunostimulation than poly I:C. This result is consistent with its enhanced effect of increasing the expression of activation markers on the DC surface (**Figure 4**).

### Comb-structured mRNA potentiates lipoplex-based mRNA vaccines against cancer

To evaluate the potential of comb-structured mRNA in cancer vaccination, we utilized anionic lipoplex, comparable with that used in clinical trials (24). Its intravenous (*i*.*v*.) injection resulted in strong protein expression from mRNA in immune tissues such as the spleen, thereby providing efficient vaccination effects (23). Herein, anionic lipoplexes were prepared by mixing cationic liposome and mRNA possessing 0, 1, 3, and 5 teeth, with an excess mRNA in charge ratio to exert negatively-charged properties. Regardless of the number of teeth, the lipoplexes showed an average size of around 200 nm with a polydispersity index below 0.12 in dynamic light scattering (DLS) measurement (**Supplementary Figure 3A**). ζ-potential of these lipoplexes was between −40 – −70 mV (**Supplementary Figure 3B**).

In this study, a lower mRNA dose (5 μg/mouse) was used compared to that used in a previous preclinical study of the comparable lipoplex (20 or 40 μg/mouse) (23), expecting that adjuvant functionalities of the comb-structured mRNA may reduce effective mRNA dose. In a reporter assay using comb-structured *firefly luciferase* (*fLuc*) mRNA, mRNA with 1 and 5 teeth exhibited fLuc expression efficiency in the spleen at a comparable level with mRNA without tooth at 24 h after injection (**Figure 6A**). Flow cytometry analyses revealed that the expression level of CD86 in splenic CD11c-positive DCs tended to increase with an increase in tooth number, with statistical significance observed between mRNA without tooth and comb-structured mRNA with 5 teeth (**Figure 6B**). This result demonstrates the appreciable potential of comb-structured mRNA for DC activation *in vivo*.

**Figure 6.**
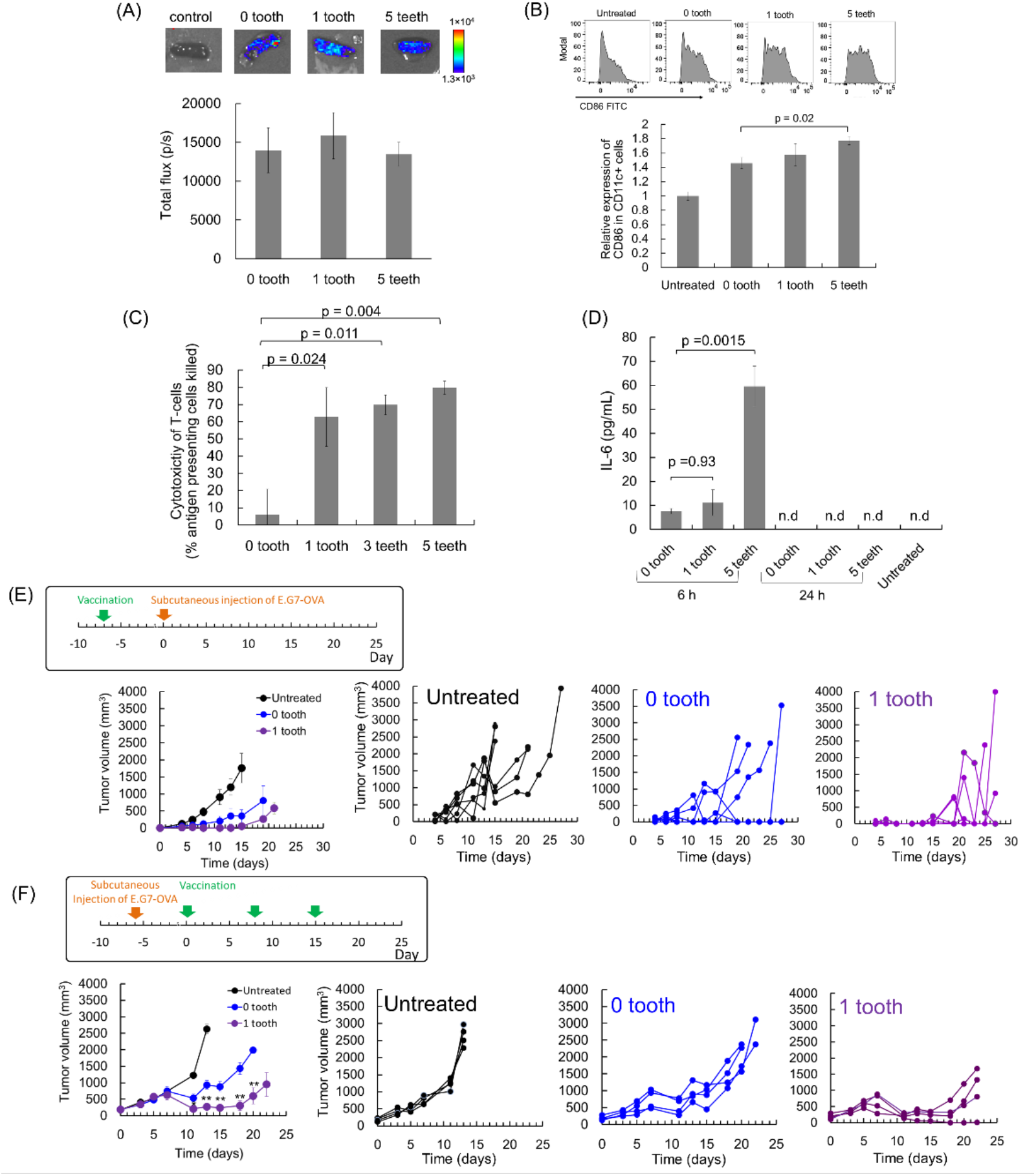
Cancer vaccines using lipoplex in mice. (**A**) Expression of fLuc in the spleen 24 h after *i*.*v*. injection of lipoplex. *n* = 4. (**B**) Activation of DC in the spleen 24 h after *i*.*v*. injection of lipoplex. *n* = 4. The expression level of CD86 in CD11c positive splenocytes. (**C**) CTL immunity against OVA 7 d after mRNA vaccination, evaluated by *in vivo* CTL assay. *n* = 4. (**D**) Serum levels of IL-6 were measured using ELISA 6 h and 24 h after *i*.*v*. injection of lipoplexes. *n* = 4. n.d.: not detected. (**E**) Prophylactic model of subcutaneously inoculated lymphoma expressing OVA, treated using *OVA* mRNA. *n* = 6 for untreated, *n* = 4 for the other two groups. (**F**) Therapeutic model of subcutaneously inoculated lymphoma expressing OVA, treated using *OVA* mRNA. *n* = 4. The average tumor volumes in each group (left) and tumor volume of individual mice (right) are shown in (**D, E**).

Vaccination effects were then assessed using mRNA expressing ovalbumin (OVA) as a model antigen by *in vivo* CTL assay, which quantifies the killing of transplanted syngeneic splenic cells presenting OVA antigen in vaccinated mice. Seven days after vaccination, comb-structured mRNA with 1, 3, and 5 teeth showed enhanced antigen-specific CTL response compared to mRNA without tooth (**Figure 6C**). The improved vaccination effect of comb-structured mRNA may be attributed to enhanced activation of DCs in immune tissues (**Figure 6B**). Enhanced secretion of proinflammatory molecules (**Figure 5**) may also contribute positively to the vaccination effect in comb-structured mRNA. In safety evaluation, we measured serum levels of proinflammatory cytokines. Introducing one tooth to mRNA does not increase the serum level of IL-6 compared to mRNA without a tooth 6 h after vaccination, while mRNA with 5 teeth showed higher IL-6 levels than those in these two groups (**Figure 6D**). Notably, the increase in IL-6 was modest in every group, and the levels returned to normal 24 h after vaccination. Serum levels of IFN-β 6 and 24 h after injection of all lipoplexes were below the detectable limit. Based on these efficiency and safety analyses, we preferentially used mRNA with one tooth throughout the following experiments to minimize safety concerns. Meanwhile, when strong adjuvant effects were necessary, mRNA with 5 teeth was also used.

Then, the therapeutic potential of comb-structured mRNA was tested firstly using a mouse model bearing subcutaneous lymphoma expressing OVA model antigen (E.G7-OVA). Mice were vaccinated before tumor inoculation in a prophylactic model and after the tumor grew to a specific size in a treatment model. In a prophylactic model, comb-structured mRNA and mRNA without a tooth suppressed tumor growth more efficiently than untreated control (**Figure 6E**). Although comb-structured mRNA tends to be more effective than mRNA without a tooth, the difference between these groups is not statistically significant due to the effective prevention of tumor growth observed in mRNA without a tooth, which led us to use a more challenging treatment model. After three-times vaccination in this treatment model, comb-structured mRNA with a tooth significantly reduced tumor volume compared to mRNA without comb-structure (**Figure 6F**).

### Comb-structured mRNA is versatile to potentiate other two types of mRNA vaccines

Recently, many studies employed iLNP via local injection for cancer vaccination (11, 13, 15), motivating us to evaluate the functionalities of comb-structured mRNA in this system. We encapsulated OVA mRNA with 0 and 1 tooth into iLNP used in an approved COVID-19 vaccine (BNT162b2), which efficiently induced cellular immunity in addition to humoral immunity (39).

Vaccination was performed via an intramuscular (*i*.*m*.) route. Notably, the size was comparable between mRNA with 0 and 1 tooth (**Supplementary Table S3**). Then, anti-OVA cellular immunity was evaluated by *in vivo* CTL assay one week after the vaccination. Among 4 tested doses of mRNA, mRNA with one tooth induced significantly enhanced antigen-specific CTL response compared to mRNA without a tooth, especially at low doses, showing substantial vaccination effects even at the mRNA dose of 0.01 μg (**Figure 7A**).

**Figure 7.**
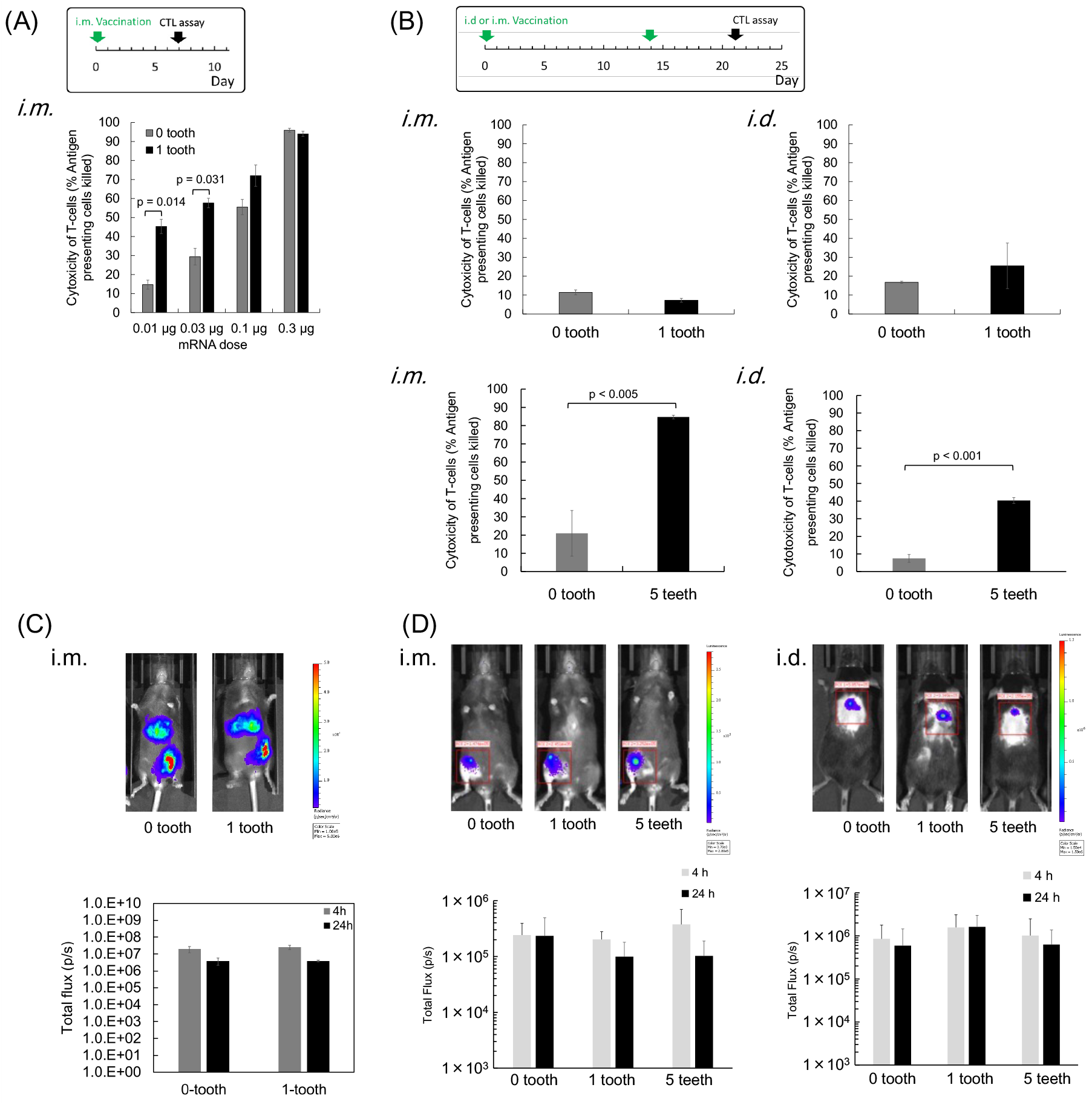
Vaccines using ionizable lipid-based LNPs (iLNPs) and polyplex micelles (PMs). (**A, B**) *In vivo* CTL assay was performed at indicated schedules. iLNPs loading *OVA* mRNA with 0 or 1 tooth were injected into mice via intramuscular (*i*.*m*.) route. *n* = 4. (**B**) PMs loading *OVA* mRNA mRNA with 0, 1, or 5 teeth were injected into mice via *i*.*m*. and intradermal (*i*.*d*.) routes. *n* = 4. (**C, D**) Luciferase expression was evaluated after *i*.*m*. injection of iLNP (**C**) and after *i*.*m*. and *i*.*d*. injection of PMs (**D**). Only expression at the muscle was quantified in (**C**). Representative images at 4 h are shown. *n* = 5 in (**C**) and *n* = 3 in (**D**).

Polymer-based systems have been rarely reported for mRNA vaccines despite their vast potential in mRNA delivery, which may be partially attributed to the low immunostimulatory properties of polymers (40). Thus, increasing the mRNA adjuvant effect using comb-structured mRNA may open the opportunity to use polymer-based systems in vaccination. In this study, we used polyplex micelle (PM), which provides efficient protein expression after *in vivo* local delivery with successful treatment outcomes in several disease models (26). Block copolymer composed of poly(ethylene glycol) (PEG) and polycation (poly(*N*-[*N*’-(2-aminoethyl)-2-aminoethyl]aspartamide): PAsp(DET)) segments was used to prepare PM by mixing mRNA and this block copolymer in an aqueous solution. Prepared PM possesses a core-shell structure with mRNA in the core surrounded by a PEG shell. In a vaccination experiment using *OVA* mRNA, PMs were injected twice at a two-week interval via *i*.*m*. and intradermal (*i*.*d*.) injection. Notably, the size was comparable between mRNA with 0 and 1 tooth (**Supplementary Table S4**). *In vivo* CTL assay performed one week after the second injection showed that mRNA with and without a tooth induced only minimal vaccination effect. There was no significant difference in the effect between these two groups (**Figure 7B**). Considering that this result is attributed to almost non-immunogenic properties of PMs (26, 41), we increased the number of teeth from 1 to 5. The increase did not largely influence PM size (**Supplementary Table S4**). In both administration routes, mRNA with 5 teeth successfully enhanced the vaccination effect for inducing CTL response compared to mRNA without a tooth (**Figure 7B**). Meanwhile, local administration of PM loading mRNA with 5 teeth did not significantly increase IL-6 and IFN-β (**Supplementary Figure S4**). Worth to notice is that tethering teeth showed only modest effects on mRNA introduction efficiency in iLNPs (**Figure 7C**) and PMs (**Figure 7D**), which may contribute to enhanced vaccination effects after the teeth tethering in these systems (**Figures 7A, B**).

## Discussion

Loading antigen and adjuvant together into a single vaccine formulation ensures co-delivery of these two components into the same antigen-presenting cells, showing promises in vaccination (8-10). Here, in alternative to an orthodox approach of integrating adjuvancy into formulating materials, we newly developed a versatile immunostimulatory comb-structured mRNA to integrate adjuvancy in mRNA cancer vaccines prepared from various existing formulation materials. Notably, the comb-structured mRNA successfully potentiated three types of existing mRNA vaccine formulations possessing completely different immunogenic properties. iLNPs intrinsically have intense adjuvant functionalities (16-18), synergizing with comb-structured mRNA adjuvant to further improve vaccination effects (**Figure 7A**). As a result, the effective dose of iLNP vaccines was able to be reduced. Meanwhile, comb-structured mRNA improved the vaccine effects of less immunostimulating anionic lipoplex (18) (**Figure 6**) and even almost non-inflammatory PM (26, 41) (**Figure 7B**). Thus, our strategy of integrating adjuvancy into comb-structured mRNA provides more freedom in choosing formulation materials for mRNA vaccines with focusing on essential properties other than immunostimulation, including the reduction of side reactions and the improvement in targetability of antigen-presenting cells.

Interestingly, optimal tooth number is dependent on the delivery formulations and routes. For example, a single tooth tethered to a small dose of mRNA (0.01 μg/mouse at minimum) was effective enough in iLNP formulation, presumably because of the intrinsic immunostimulatory properties of the iLNP. In contrast, 5 teeth-tethered mRNA at an increased dose (12, 20 μg/mouse) for the PM formulation was optimum for effective vaccination without a systemic increase in inflammatory molecules. Furthermore, one tooth at mRNA dose of 5 μg/mouse was optimal for anionic lipoplex to exert an efficient antitumor effect without significantly increasing serum cytokine levels. These results highlight that the immunostimulation intensity of comb-structured mRNA systems is systematically adjusted to various mRNA vaccine formulations. Comb-structured mRNA does not require additional exogenous materials for immunostimulation other than RNA composed of naturally existing nucleotides. This characteristic is a potential advantage of safety in clinical application. Remarkably, tethering of only one short dsRNA tooth (24 bp) to a long mRNA strand drastically induced the immunological response with a negligible increase in total RNA doses, as demonstrated in the immunostimulation effect of 783 nt *gLuc* mRNA (**Figures 2-5**) and the vaccination effect of 1437 nt *OVA* mRNA (**Figures 6, 7**). Such short teeth introduction minimally affected physicochemical properties of mRNA formulations, as shown in **Supplementary Figure S3** for lipoplex, **Supplementary Table S3** for iLNP, and **Supplementary Table S4** for PM, respectively, indicating the broad applicability of this approach in various types of mRNA formulations.

Simple rationality in adjuvant design is also merit for this dsRNA tooth-based approach. Fine-tuning the structure of just a short dsRNA tooth (24 bp) allows sufficient adjuvant functionalities. Interestingly, subtle changes in the dsRNA structure largely influence the immunostimulation intensity (**Figure 2**). Although detailed insights into the mechanisms underlying this phenomenon are yet to be clarified, it would be worth discussing the possible scheme of RIG-I recognition. According to previous reports, the 5’ triphosphate structure at the blunt end is a binding motif of RIG-I recognition, and a slight disturbance of the blunt structure, *i*.*e*., the presence of 5’ or 3’ overhang with a few base-pairs, drastically decreases immune activation (22, 27). In addition, RIG-I requires at least 20 nt dsRNA structure for its activation. Thus, unintended hybridization products, including loop structure or dsRNA with 5’ or 3’ overhang, might decrease the RIG-I recognition, depending on tooth dsRNA sequence and length. Another possible determinant in RIG-I recognition is the dsRNA microenvironment. Indeed, the gap sequence length between dsRNA tooth and mRNA and the binding of a tooth to mRNA largely influenced immunostimulation with a tooth with 2 nt gap sequence inducing the expression of inflammatory molecules more efficiently than that with 10 nt gap sequence and without attaching to mRNA (**Figures 2, 5, S2**). Among these teeth, a tooth with 2 nt gap sequence locates closest to the 17 nt dsRNA from cRNA and mRNA along mRNA strands. The 17 nt cRNA/mRNA dsRNA might facilitate the binding of the nearby tooth to RIG-I or be recognized simultaneously with the tooth by RIG-I, although comprehensive mechanistic analyses are needed to examine this hypothesis. Practically, even if dsRNA teeth were unexpectedly detached from mRNA after administration into the body, the detached teeth would induce minimal off-target inflammation in cells and tissues that do not show antigen expression from mRNA.

Previously, we reported the preparation of immunostimulatory mRNA by hybridizing poly U sequence in its polyA tail region, which is the first attempt to integrate adjuvancy directly into antigen-encoding mRNA for vaccine formulation (42). The approach in the present study has several advantages over this previous approach. First, only the present comb-structured mRNA can control the immunostimulation intensity, providing advantages in safety and versatility as aforementioned. Second, the current comb-structured approach uses chemically synthesized cRNA for precise preparation of the blunt-ended dsRNA with 5’ triphosphate, a binding motif for RIG-I activation. In contrast, preparation of such a structure was unachievable in the previous approach, which entirely relies on IVT for poly U preparation. More critically, IVT preparation of poly U in the previous approach may produce contaminant RNA strands complementary to poly U via an uncontrolled process of RNA-templated RNA transcription during IVT (28-30), and dsRNA from poly U and the contaminant RNA may influence the immunostimulating activity of mRNA hybridized with poly U. The current approach circumvents the formation of such by-products by justifying the sequences of 5’ppp-RNA in combination with the proper chemical synthesis of cRNA.

The present system drastically increased the secretion of IFN-β *in vitro*, although the influence of type I interferon on mRNA vaccines is still controversial. Previous studies showed either positive or negative effects of type I interferon on cellular immunity induction by mRNA vaccines (23, 43-46). Although the precise mechanisms underlying positive outcomes of comb-structured mRNA in the present study need further investigation, comb-structured mRNA induced the secretion of various proinflammatory molecules (**Figure 5**). Accordingly, some of the molecules might influence positively to override the negative effect of type I interferon. Notably, comb-structured mRNA controls the intensity of type I interferon induction, potentially helping avoid its negative influences.

In conclusion, our system is a simple and robust platform to supply a desired intensity of immunostimulation to various mRNA cancer vaccine systems and improve their efficiency only by adding a small amount of RNA. While the current study focuses on system development and fundamental analyses, future studies are planned to optimize the system using models more relevant to the clinical settings from the standpoint of clinical translation.

## Materials and Methods

### Materials

*fLuc* and *OVA* mRNA were purchased from TriLink® Biotechnologies (San Diego, CA). mRNA encoding *gLuc* and *gp100* was prepared by IVT using mMessage mMachine T7 kit (Thermofisher, Waltham, MA). DNA templates are prepared by inserting protein-coding sequence of *gLuc* (pCMV-GLuc control vector, New England BioLabs, Ipswich, MA) into pSP73 vectors (Promega, Madison, WI) possessing 120 bp polyA/T sequences downstream of a multiple cloning site. cRNA was purchased from Hokkaido System Bioscience (Hokkaido, Japan) (**Supplementary Table S2**), 5’ppp-RNA was prepared by IVT in the absence of ATP using MEGAshortscript T7 kit (Thermofisher) and DNA template, with sequences listed in **Supplementary Table S1**. HEPES (4-(2-hydroxyethyl)-1-piperazineethanesulfonic acid) 1 M was purchased from Gibco (Waltham, MA), sodium chloride 5 M was purchased from Thermo Fisher Scientific, while chloroform was purchased from Wako Pure Chemicals Industries (Osaka, Japan). DOTMA (1,2-di-O-octadecenyl-3-trimethylammonium propane (chloride salt)) and DOPE (1,2-dioleoyl-sn-glycero-3-phosphoethanolamine) were purchased from NOF Corporation (Tokyo, Japan), while Lipofectamine LTX Plus was purchased from Thermo Fisher Scientific. DC2.4 Mouse Dendritic Cell was purchased from MilliporeSigma (Burlington, MA). RAW-Lucia™ ISG (wild type), RAW-Lucia™ ISG-KO-RIG-I, RAW-Lucia™ ISG-KO-MDA5, HEK-Blue™ Null, HEK-Blue™ hTLR3, and HEK-Blue™ TLR7 were purchased from Invivogen (San Diego, CA). E.G7-OVA and B16F0-Luc were obtained from JCRB Cell Bank (Tokyo, Japan). Normal Human Dendritic Cell was purchased from Lonza Group AG (Basel, Switzerland). RPMI-1640 (Fujifilm Wako Chemicals, Osaka, Japan), DMEM (Sigma-Aldrich, St. Louis, MO), GM-3TM Lymphocyte Growth Medium-3 (LGM-3™) (Lonza), Opti-MEM Serum-Free medium (Thermofisher), PBS (Fujifilm Wako Chemicals, Osaka, Japan), FBS (Dainippon Sumitomo Pharma Co. Ltd., Osaka, Japan), penicillin-streptomycin (Sigma-Aldrich), normocin (Invivogen), blasticidin (Invivogen), Geneticin selective antibiotic G418 Sulfate (Gibco), granulocyte-macrophage colony-stimulating factor (GM-CSF) 200-15 (Shenandoah BioTech, Warminster, PA). Mercaptoethanol (Thermofisher), Glutamax™ (Gibco), sodium pyruvate 100 mmol/L (Fujifilm), recombinant human granulocyte-macrophage colony-stimulating factor (GM-CSF) (R&D Systems, Minneapolis, MN), recombinant human interleukin 4 (IL-4) (R&D Systems), Alpha MEM with nucleosides (StemCell Technologies, Vancouver, Canada) were purchased for cell culture purposes. PE-conjugated anti-mouse CD40 (eBioScience™ 12-0401-81), CD86 (eBioScience™ 12-0862-81), MHC Class I (H-2Kb) Monoclonal Antibody (eBioscience™ 12-5958-82), MHC Class II (I-A) Monoclonal Antibody (NIMR-4) (eBioscience™ 12-5322-81) were purchased from Thermo Fisher Scientific. TruStain FcX™ anti-mouse CD16/32 antibody (Biolegend 101319), FITC anti-mouse CD86 antibody (Biolegend 105005), APC anti-mouse CD11c antibody (Biolegend 117309), human TruStain FcX™ (Biolegend 422301), and PE anti-human CD86 antibody (Biolegend 305405) were purchased from Biolegend (San Diego, CA). Cell staining buffer (BioLegend 420201), Accutase (Sigma-Aldrich), ACK lysing buffer (Thermofisher), CellTrace™ CFSE Cell Proliferation Kit (Thermofisher), luciferin (Summit pharmaceuticals, Tokyo, Japan), IVT RNA kit, SIINFEKL (Thermofisher), Renilla Luciferase Assay System (Promega) Quanti-Luc (Invivogen), Secreted Embryonic Alkaline Phosphatase (SEAP) (Invivogen) were purchased for various cell processing and analysis experiments. Taqman primer/probe mixtures (Applied Biosystems) for mouse IL6 (FAM, Mm00446190), mouse INFB1 (FAM, Mm00439552_s1) and Mouse ACTB (Actin, Beta) Endogenous Control (FAM™ Dye/MGB probe, Non-Primer Limited) (Thermofisher) and Applied biosystems Taqman™ Fast Universal PCR Master Mix (2x) No Amperase™ (Applied Biosystem) were purchased for quantitative PCR experiments. Mouse IL6 Duoset ELISA kit (R&D systems) and Mouse IFN-β Duoset ELISA kit (R&D systems) were purchased for ELISA experiments. C57BL/6J mice (7 weeks, female) were purchased from Oriental Kobo.

### Preparation of comb-structured mRNA

Comb-structured mRNAs were prepared from 10 mM HEPES buffered solution of mRNA, cRNA, and 5’ppp-RNA composed at appropriate molar equivalents. The solution was heated at 65°C for 5 minutes using Takara® PCR Thermal Cycler (Takara Corp., Shiga, Japan), and then gradually cooled down to 30°C over a time span of 10 minutes before settling down to 4 °C. Successful hybridization of dsRNA teeth to mRNA was confirmed by quantifying unreacted Cy5-labeled dsRNA in the flow-through fraction after ultrafiltration. Namely, comb-structured mRNA composed of *OVA* mRNA, cRNA, and Cy5-labeled 5’ppp-RNA was prepared in the solution. Then, the solution was subjected to ultrafiltration using an ultra-0.5 centrifugal filter of MWCO 100K (Merck KGaA, Darmstadt, Germany) under centrifugation at 4 °C for 5 minutes at 10,000 G. The emission at 670 nm of the flow-through was measured using a TECAN plate reader (emission 651 nm) to detect Cy5.

### Preparation of anionically-charged mRNA lipoplex

Anionically-charged mRNA lipoplex was prepared according to the previous report (23). First, a cationic liposome was prepared from DOTMA and DOPE (1:1 molar ratio, 6 mM total lipid concentration) using a thin-film hydration and sonication method. Next, mRNA or comb-structured mRNA was dissolved in water containing NaCl and mixed with the liposome suspension at a nitrogen/phosphate (N/P) ratio of 0.5 to obtain anionically-charged mRNA lipoplex with a final NaCl concentration of 150 mM.

### Preparation of Lipofectamine LTX with Plus reagent/mRNA lipoplex

Lipofectamine LTX with Plus reagent/mRNA lipoplex was prepared according to the manufacturer’s protocol. 3.5 μg of mRNA was dissolved in 350 μL of Opti-MEM^®^ Reduced Serum Medium containing 3.5 μL of Plus™ reagent. This mRNA solution was then added to a 350 μL of Opti-MEM^®^ Reduced Serum Medium containing 8.75 μL of lipofectamine^®^ LTX reagent. Lipofectamine LTX with Plus reagent/comb-structured mRNA lipoplexes were prepared similarly, with the total RNA amount including mRNA, cRNA, and 5’ppp-RNA used for determining mixture ratio of RNA with lipofectamine^®^ LTX and Plus™ reagents.

### Preparation of ionizable lipid-based LNP

Ionizable lipid, ALC-0315 (MedChemExpress, Monmouth Junction, NJ, USA), phospholipid, 1,2-distearoyl-sn-glycero-3-phosphocholine (DSPC, Fujifilm Wako, Osaka, Japan), cholesterol (Sigma Aldrich), and PEG-lipid, ALC-0159 (MedChemExpress) were dissolved in ethanol at the molar ratio of 46.3: 9.4: 42.7: 1.6 in molar ratio (total lipid concentration: 35 mM). mRNA was dissolved in 50 mM citrate buffer (pH 3) at the concentration of 459 μg/mL. mRNA and lipid solution were mixed at volume ratio of 2:1 using microfluidics (NanoAssemblr® Spark, Precision Nanosystems, Vancouver, Canada) to prepare LNP at N/P ratio of 6, followed by 40-fold dilution using PBS and ultrafiltration using Amicon Ultra-15-30K centrifugal units (Merck Millipore) for buffer replacement.

### Preparation of mRNA polyplex micelles

A block copolymer of poly(ethylene glycol) (Mw: 44 kDa) with poly(N-[N’-(2-aminoethyl)-2-aminoethyl]aspartamide) [PEG-PAsp(DET)] was synthesized as previously reported (47). The polymerization degree of the PAsp(DET) segment was determined to be 59 from the ^1^H-NMR (400 MHz, JEOL ECS-400, JEOL, Tokyo, Japan) measurement. PEG-PAsp(DET) was dissolved in 10 mM HEPES buffer. This solution was mixed with mRNA in the same buffer at an N/P ratio of 5 to obtain polyplex micelles (PM). The final concentration of mRNA or comb-structured mRNA was fixed to 300 μg/mL.

### Hydrodynamic diameter (D_h_) and Zeta potential determination by Zetasizer

DLS measurement was conducted by Zetasizer Nanoseries (Malvern Instruments Ltd., Malvern, UK) to measure D_h_ of lipoplex and polyplex micelle formulations of *OVA* mRNA with and without dsRNA teeth. The measurement was performed at a detection angle of 173° and a temperature of 25°C. The refractive index (RI) was set to 1.45 (RI of phospholipid) for lipoplex and ionizable lipid-based LNP and 1.59 (RI of polystyrene) for polyplex micelle, respectively. The D_h_ was obtained by treating the decay rate in the photon correlation function by the cumulant method and applying the Stokes-Einstein equation. The zeta potential was determined by laser-doppler electrophoresis using the same instrument. According to the obtained electrophoretic mobility, the zeta potential of each sample was calculated according to the Smoluchowski equation: ζ = (4πημ)/ε, where η is the viscosity of the solvent, μ is the electrophoretic mobility, and ε is the dielectric constant of the solvent.

### Evaluation of proinflammatory cytokine transcripts induced by comb-structured mRNA in DC2.4 cells

DC2.4 cells were seeded onto 24-well plates at a 2 × 10^5^ cells/well density. 24 h later, the culture medium was replaced with serum-free Opti-MEM medium (500 μL) and added with 0.5 μg of *gLuc* mRNA with or without teeth formulated with Lipofectamine LTX and Plus reagents. After 4 h of incubation, cellular mRNA was extracted using RNeasy^®^ Mini Kit (Qiagen N.V., Düsseldorf, Germany), converted to complementary DNA (cDNA) using ReverTra Ace™ qPCR RT Master Mix with gDNA Remover (Toyobo Inc., Osaka, Japan) for quantitative real-time PCR.

### Probing of RIG-I and MDA-5 involvement to immunogenicity using RAW-Lucia™ ISG Cells (IFN Reporter RAW 264.7 macrophages)

*OVA* mRNA (100 ng) with or without teeth formulated with Lipofectamine LTX and Plus reagents in 20 μL Opti-MEM medium was added to each well of 96 plates. Then, RAW-Lucia ISG cells (100,000 cells/well) suspended in 180 mL of DMEM containing 10% FBS and 100 μg/mL PS were added to each well. Of note, RAW-Lucia™ ISG (wild type), RAW-Lucia™ ISG-KO-RIG-I, and RAW-Lucia™ ISG-KO-MDA5 cells were cultured in DMEM containing 10% FBS, 100 μg/mL PS, and 100 μg/mL normocin prior to maintain genetic stability of these transgenic cell lines. After 4 h, the medium was collected to measure lLuc expression.

### Probing of hTLR3 and TLR7 involvement in immunogenicity using hTLR3-Blue-HEK and TLR7-Blue-HEK293

HEK-Blue™ Null, HEK-Blue™ hTLR3, and HEK-Blue™ TLR7 cells were cultured in a selection medium according to the manufacturer’s protocol. *gLuc* mRNA (100 ng) with or without teeth formulated with Lipofectamine LTX and Plus reagents in 20 μL Opti-MEM medium was added to each well of 96 plates. HEK-Blue™ hTLR3 cells (500,000 cells/well) or HEK-Blue™ TLR7 (400,000 cells/well) and HEK-Blue™ Null (500,000 cells/well) were suspended in 180 μL of SEAP medium, respectively, and were added to each well. After 16 h of incubation, the plates were examined for absorbance at 630 nm using a TECAN microplate reader. Relative absorbance values are shown in **Figure 3B**.

### Gene expression, surface antigen presentation, and cytokines production in primary BMDCs

Primary BMDCs were obtained according to a previous report (48). In brief, the cells were flushed out from the tibia and femur bone marrow of C57BL/6J. The cells were cultured at 37°C in 5% CO_2_ in an R10 culture medium containing granulocyte-macrophage colony-stimulating factor (GM-CSF). On day 8, the floating cells in the medium were collected. 400,000 cells suspended in 2 mL of R10 medium was added to each well of a 12-well plate and transfected on the same day with 1 μg/well of *gLuc* mRNA formulated with Lipofectamine LTX and Plus reagents. 10 μL of the medium at several time points was collected for gLuc expression assay using Renilla Luciferase Assay System (Promega) and a luminometer, GloMax-Multi+ Detection System (Promega). After 24 h, both floating and attaching cells were collected, washed, blocked with anti-mouse CD16/31 antibody (1/100 dilution in cell staining buffer), and stained with PE-conjugated antibody mouse CD40 (0.005 μg/mL), CD86 (0.005 μg/mL), MHCI (0.007 μg/mL) and MHCII (0.003 μg/mL) for 20 min on ice. The immunostained cells were then washed, stained with DAPI, and examined by flow cytometry (Cell Analyzer LSRFortessa X-20, Becton Dickinson).

In another batch of the experiment, the medium from the transfected cells was collected to quantify the expression of immune molecules at protein levels. Luminex® Multiplex Assays (ThermoFisher Scientific) was performed to quantify expression levels of 24 molecules, including G-CSF, IFN-γ, IL-1α, 1β, 2, 4, 5, 6, 7, 9, 10, 12p40, 12p70, 13, 15, 17, TNF-α, CXCL1, 2, 10, CCL2, 3, 4, 5. Expression of IFN-*β* was quantified using Mouse IFN-*β* Duoset ELISA kit.

### Human DC

Human dendritic cell purchased from Lonza was seeded to 12-well plates at 160,000 cells/well containing 2 mL of medium. Cells were transfected with mRNA (0.5 μg/well) using Lipofectamine LTX and Plus reagent. Adherent and non-adherent cells were harvested after 24h for immunostaining. Adherent cells were detached from the plate using 400 μL of accutase solution for 5 min, and the reaction was stopped using a culture medium. All cells were centrifuged at 500 g for 5 min at 4°C. Cells were blocked with 100 μL of blocking trustain human antibody (1/250 dilution in cell staining buffer) and kept on ice for 10 min. Then, cells were stained with 100 μL of human PE-CD86 solution (BioLegend) (5 μg/mL) and kept on ice for 20 minutes. After two rounds of washing using 500 μL of cell staining buffer, cells were reconstituted with 500 μL of staining buffer. The cells were analyzed by flow cytometry using BD LSRFortessa™ X-20.

### Expression of luciferase mRNA in mice

C57BL/6J mice were intravenously injected with 5 μg/mouse of *fLuc* mRNA formulated with lipoplex. After 24 h, the spleen was extracted for measuring fLuc expression using IVIS Spectrum SP-BFM-T1 (Perkin Elmer) after an intraperitoneal injection of 3 mg of luciferin. fLuc IVIS experiment was also conducted for mice injected with ionizable lipid based LNPs via *i*.*m*. route, and mRNA PMs *via i*.*d*. and *i*.*m* routes (1 μg mRNA/mouse). All animal experiments were conducted under the approval of the animal care and use committees in the Innovation Center of NanoMedicine, Kawasaki Institute of Industrial Promotion (Kanagawa, Japan), and Kyoto Prefectural University of Medicine (Kyoto, Japan).

### Examination of surface antigen presentation of dendritic cells in vivo

C57BL/6J mice (7 weeks, female) were intravenously injected with 5 μg of *OVA* mRNA/mouse formulated with lipoplex. After 24 h, splenocytes were extracted from spleens of donor C57BL6 mice and added with ACK lysing buffer to destroy the red blood cells. 5 × 10^6^ cells were taken and centrifuged at 400 G for 5 min at 4 °C, and then resuspended in cell staining buffer. The cells were later blocked with 10 μg/mL CD16/32 antibody in 100 μL solution and kept on ice for 15 min. Thereafter, the cells were stained with a 100 μL mixture of FITC-CD86 (10 μg/mL) and APC-CD11c (25 μg/mL) antibody in cell staining buffer for 20 min on ice. The cells were washed 3 times and then resuspended in 1 mL of cell staining buffer for examination using flow cytometry.

### In vivo cytotoxic T lymphocyte (CTL) assay

C57BL/6J mice were vaccinated by intravenous (*i*.*v*.) injection of 5 μg of *OVA* mRNA/mouse formulated with lipoplex. After 7 days, *in vivo* CTL assay was performed by following a previous protocol (49). In brief, splenocytes were extracted from donor C57BL6 mice and added with ACK lysing buffer to destroy the red blood cells. The remaining cells were washed and divided into two groups. One group was treated with OVA257-264 peptide (SIINFEKL, Toray, Tokyo, Japan) for 1 h in an incubator (37 °C, CO_2_ 5%), and later dyed with 5 μM CFSE (CellTrace™ CFSE Cell Proliferation Kit - For Flow Cytometry, Molecular Probes, Eugene, OR) (1 mL PBS containing CFSE 5 μM per 3 × 10^7^ splenic cells) for 10 min in a water bath at 37 °C. The other group was left non-treated, incubated for 1 h (37 °C, CO_2_ 5%), and later dyed with 0.5 μM CFSE (1 mL PBS containing CFSE 5 μM per 3 × 10^7^ splenocytes) for 10 min in a water bath at 37 °C. Both groups were concentrated to 5 × 10^7^ splenocytes per 1 mL PBS and placed on an ice bath before mixing prior to the injection into mice. The splenocytes (1 × 10^7^ cells in 200 μL of suspension) were intravenously injected into vaccinated mice. One day later, the vaccinated mice were sacrificed to check the viability of the injected CFSE-labeled donor splenocytes by flow cytometry (BD LSRFortessa™ X-20).

For iLNP vaccines, C57BL6 mice were vaccinated via *i*.*m*. injection of 100 μL iLNP solution containing indicated mRNA amount. Seven days after the vaccination, *in vivo* CTL assay as performed as described above.

For mRNA-PM vaccines, C57BL6 mice received the vaccine twice at the interval of 2 weeks with *OVA* mRNA dose set to be 20 μg/mouse for *i*.*m*. vaccination, and 12 μg/mouse for *i*.*d*. vaccination, in each of the prime and boost vaccination. In the *i*.*m*. vaccination, 20 μg mRNA formulated in PM in 100 μL solution was divided into two and injected into the thigh muscle on both left and right sides. In *i*.*d*. vaccination, 12 μg mRNA formulated in PM in 50 μL solution was divided into five and injected at 5 locations in the dermis of the back of the mouse. On day 6, post-booster dose, mice were subjected to *in vivo* cytotoxicity assay as described above.

### Prophylaxis tumor suppression study against subcutaneously implanted lymphoma model

C57BL/6J mice were vaccinated one time with *i*.*v*. injection of 5 μg of *OVA* mRNA with or without teeth formulated with anionic lipoplex. After 7 days, mice were subcutaneously implanted with 8 × 10^5^ E.G7-OVA cells. The tumor growth was followed for 2 months. The tumor volume was calculated by the formula (1/2 × long axis × (short axis)^2^).

### Therapeutic tumor suppression study against subcutaneously implanted lymphoma model

C57BL/6J mice were subcutaneously implanted with 1 × 10^6^ E.G7-OVA cells. The therapeutic intervention was started 6 days after the inoculation when the tumor grew to an average size of 185 mm^3^ for each group. On this day, and 8 and 15 days later, mice were vaccinated with *i*.*v*. injection of 5 μg of *OVA* mRNA with or without teeth formulated with anionic lipoplex. The tumor growth was followed as described in the previous section.

### Safety assay by examining IL-6 and IFN-β content in serum

C57BL/6J mice were intravenously injected with 5 μg mRNA with or without teeth formulated in anionic lipoplex. After 6 and 24 hours, mice were sacrificed to obtain serum. The serum was examined for IL-6 and IFN-β content using ELISA.

### Data presentation

Data are presented as mean ± standard error of the mean (SEM). In **Figures 6B,F and 7**, statistical analyses are performed using unpaired two-tailed Student’s t-test. In **Figures 2, 3, 4, 6C,D** statistical analyses are performed using analysis of variance (ANOVA) followed by Tukey’s post hoc test.

## Acknowledgments

This work was supported by the Center of Innovation Program (COI) from Japan Science and Technology Agency (JST), Grants-in-Aid for Challenging Research (Pioneering) [18H05378 to K.K.], for Scientific Research (A) [21H04962 to S.U., 21H04967 to K.K.] and (B) [18K03529 to S.U.] from the Ministry of Education, Culture, Sports, Science and Technology, Japan (MEXT), Leading Advanced Projects for Medical Innovation [21gm0010008s0101 to S.U.], and Research Program on Emerging and Re-emerging Infectious Diseases [21fk0108620h0001 to S.U.] from Japan Agency for Medical Research and Development (AMED). We would like to thank Hiroaki Kinoh, Xueying Liu, Joachim Van Guyse, Keisuke Nagao, Hikaru Saitoh, Yuki Sato, Yuki Tada (iCONM), Satomi Nakagahara (NanoCarrier Ltd.), for their technical assistance.

## Author Contributions

Conceptualization: SU, Methodology: TAT, SA, MM, NY, EB, ZW, SF, Investigation: TAT, SA, MM, NY, EB, ZW, SF, Supervision: KK, SU, Writing—original draft: TAT, SU, Writing—review & editing: SA, KK, SU

## References

1. N. Pardi, M. J. Hogan, F. W. Porter, D. Weissman, mRNA vaccines - a new era in vaccinology. Nat. Rev. Drug Discov. 17, 261–279 (2018).

2. L. Miao, Y. Zhang, L. Huang, mRNA vaccine for cancer immunotherapy. Mol. Cancer 20, 41 (2021).

3. A. J. Barbier, A. Y. Jiang, P. Zhang, R. Wooster, D. G. Anderson, The clinical progress of mRNA vaccines and immunotherapies. Nature Biotechnol. 40, 840–854 (2022).

4. U. Sahin et al., Personalized RNA mutanome vaccines mobilize poly-specific therapeutic immunity against cancer. Nature 547, 222–226 (2017).

5. G. Cafri et al., mRNA vaccine–induced neoantigen-specific T cell immunity in patients with gastrointestinal cancer. J. Clin. Invest. 130, 5976–5988 (2020).

6. B. Weide et al., Direct injection of protamine-protected mRNA: results of a phase 1/2 vaccination trial in metastatic melanoma patients. J. Immunother. 32, 498–507 (2009).

7. S. Van Lint et al., Preclinical evaluation of TriMix and antigen mRNA-based antitumor therapy. Cancer Res. 72, 1661–1671 (2012).

8. E. Schlosser et al., TLR ligands and antigen need to be coencapsulated into the same biodegradable microsphere for the generation of potent cytotoxic T lymphocyte responses. Vaccine 26, 1626–1637 (2008).

9. N. O. Fischer et al., Colocalized delivery of adjuvant and antigen using nanolipoprotein particles enhances the immune response to recombinant antigens. J. Am. Chem. Soc. 135, 2044–2047 (2013).

10. R. Kuai, L. J. Ochyl, K. S. Bahjat, A. Schwendeman, J. J. Moon, Designer vaccine nanodiscs for personalized cancer immunotherapy. Nature Mater. 16, 489–496 (2017).

11. M. A. Oberli et al., Lipid Nanoparticle Assisted mRNA Delivery for Potent Cancer Immunotherapy. Nano Lett. 17, 1326–1335 (2017).

12. O. A. W. Haabeth et al., mRNA vaccination with charge-altering releasable transporters elicits human T cell responses and cures established tumors in mice. Proceedings of the National Academy of Sciences 115, E9153–E9161 (2018).

13. L. Miao et al., Delivery of mRNA vaccines with heterocyclic lipids increases antitumor efficacy by STING-mediated immune cell activation. Nature Biotechnol. 37, 1174–1185 (2019).

14. H. Zhang et al., Delivery of mRNA vaccine with a lipid-like material potentiates antitumor efficacy through Toll-like receptor 4 signaling. Proc. Natl. Acad. Sci. U. S. A. 118 (2021).

15. M. A. Islam et al., Adjuvant-pulsed mRNA vaccine nanoparticle for immunoprophylactic and therapeutic tumor suppression in mice. Biomaterials 266, 120431 (2021).

16. M. G. Alameh et al., Lipid nanoparticles enhance the efficacy of mRNA and protein subunit vaccines by inducing robust T follicular helper cell and humoral responses. Immunity 54, 2877–2892 e2877 (2021).

17. S. Ndeupen et al., The mRNA-LNP platform’s lipid nanoparticle component used in preclinical vaccine studies is highly inflammatory. iScience 24, 103479 (2021).

18. S. Tahtinen et al., IL-1 and IL-1ra are key regulators of the inflammatory response to RNA vaccines. Nat. Immunol. 23, 532–542 (2022).

19. S. Abbasi, S. Uchida, Multifunctional Immunoadjuvants for Use in Minimalist Nucleic Acid Vaccines. Pharmaceutics 13, 644 (2021).

20. S. Heidegger et al., RIG-I activating immunostimulatory RNA boosts the efficacy of anticancer vaccines and synergizes with immune checkpoint blockade. EBioMedicine 41, 146–155 (2019).

21. J. Koerner et al., PLGA-particle vaccine carrying TLR3/RIG-I ligand Riboxxim synergizes with immune checkpoint blockade for effective anti-cancer immunotherapy. Nature Commun. 12, 2935 (2021).

22. M. Schlee et al., Recognition of 5’ triphosphate by RIG-I helicase requires short blunt double-stranded RNA as contained in panhandle of negative-strand virus. Immunity 31, 25–34 (2009).

23. L. M. Kranz et al., Systemic RNA delivery to dendritic cells exploits antiviral defence for cancer immunotherapy. Nature 534, 396–401 (2016).

24. U. Sahin et al., An RNA vaccine drives immunity in checkpoint-inhibitor-treated melanoma. Nature 585, 107–112 (2020).

25. X. Hou, T. Zaks, R. Langer, Y. Dong, Lipid nanoparticles for mRNA delivery. Nature Reviews Materials 10.1038/s41578-021-00358-0 (2021).

26. S. Uchida, K. Kataoka, Design concepts of polyplex micelles for in vivo therapeutic delivery of plasmid DNA and messenger RNA. Journal of Biomedical Materials Research Part A 107, 978–990 (2019).

27. A. Schmidt et al., 5’-triphosphate RNA requires base-paired structures to activate antiviral signaling via RIG-I. Proc. Natl. Acad. Sci. U. S. A. 106, 12067–12072 (2009).

28. C. Cazenave, O. C. Uhlenbeck, RNA template-directed RNA synthesis by T7 RNA polymerase. Proc. Natl. Acad. Sci. U. S. A. 91, 6972–6976 (1994).

29. F. J. Triana-Alonso, M. Dabrowski, J. Wadzack, K. H. Nierhaus, Self-coded 3’extension of run-off transcripts produces aberrant products during in vitro transcription with T7 RNA polymerase. J. Biol. Chem. 270, 6298–6307 (1995).

30. K. Kariko, H. Muramatsu, J. Ludwig, D. Weissman, Generating the optimal mRNA for therapy: HPLC purification eliminates immune activation and improves translation of nucleoside-modified, protein-encoding mRNA. Nucleic Acids Res. 39, e142 (2011).

31. Y. Gholamalipour, A. Karunanayake Mudiyanselage, C. T. Martin, 3’ end additions by T7 RNA polymerase are RNA self-templated, distributive and diverse in character-RNA-Seq analyses. Nucleic Acids Res. 46, 9253–9263 (2018).

32. N. Yoshinaga et al., Induced packaging of mRNA into polyplex micelles by regulated hybridization with a small number of cholesteryl RNA oligonucleotides directed enhanced in vivo transfection. Biomaterials 197, 255–267 (2019).

33. H. Kato et al., Length-dependent recognition of double-stranded ribonucleic acids by retinoic acid-inducible gene-I and melanoma differentiation-associated gene 5. J. Exp. Med. 205, 1601–1610 (2008).

34. I. Botos, L. Liu, Y. Wang, D. M. Segal, D. R. Davies, The toll-like receptor 3:dsRNA signaling complex. Biochim. Biophys. Acta 1789, 667–674 (2009).

35. F. Heil et al., Species-Specific Recognition of Single-Stranded RNA via Toll-like Receptor 7 and 8. Science 303, 1526–1529 (2004).

36. C. Pollard et al., Type I IFN counteracts the induction of antigen-specific immune responses by lipid-based delivery of mRNA vaccines. Mol. Ther. 21, 251–259 (2013).

37. S. Uchida, K. Kataoka, K. Itaka, Screening of mRNA chemical modification to maximize protein expression with reduced immunogenicity. Pharmaceutics 7, 137–151 (2015).

38. M. P. Kostinov, N. K. Akhmatova, E. A. Khromova, A. M. Kostinova, Cytokine Profile in Human Peripheral Blood Mononuclear Leukocytes Exposed to Immunoadjuvant and Adjuvant-Free Vaccines Against Influenza. Front. Immunol. 11, 1351 (2020).

39. A. B. Vogel et al., BNT162b vaccines protect rhesus macaques from SARS-CoV-2. Nature 592, 283–289 (2021).

40. A. K. Blakney et al., Polymeric and lipid nanoparticles for delivery of self-amplifying RNA vaccines. J. Control. Release 338, 201–210 (2021).

41. S. Uchida et al., In vivo messenger RNA introduction into the central nervous system using polyplex nanomicelle. PLoS One 8, e56220 (2013).

42. S. Uchida et al., Designing immunostimulatory double stranded messenger RNA with maintained translational activity through hybridization with poly A sequences for effective vaccination. Biomaterials 150, 162–170 (2018).

43. K. Broos et al., Particle-mediated Intravenous Delivery of Antigen mRNA Results in Strong Antigen-specific T-cell Responses Despite the Induction of Type I Interferon. Molecular Therapy - Nucleic Acids 5, e326 (2016).

44. V. K. Udhayakumar et al., Arginine-Rich Peptide-Based mRNA Nanocomplexes Efficiently Instigate Cytotoxic T Cell Immunity Dependent on the Amphipathic Organization of the Peptide. Adv Healthc Mater 6, 1601412 (2017).

45. L. Van Hoecke et al., The Opposing Effect of Type I IFN on the T Cell Response by Non-modified mRNA-Lipoplex Vaccines Is Determined by the Route of Administration. Mol Ther Nucleic Acids 22, 373–381 (2020).

46. C. Li et al., Mechanisms of innate and adaptive immunity to the Pfizer-BioNTech BNT162b2 vaccine. Nat. Immunol. 23, 543–555 (2022).

47. N. Kanayama et al., A PEG-based biocompatible block catiomer with high buffering capacity for the construction of polyplex micelles showing efficient gene transfer toward primary cells. ChemMedChem 1, 439–444 (2006).

48. M. B. Lutz et al., An advanced culture method for generating large quantities of highly pure dendritic cells from mouse bone marrow. J. Immunol. Methods 223, 77–92 (1999).

49. Y. Yang, C. T. Huang, X. Huang, D. M. Pardoll, Persistent Toll-like receptor signals are required for reversal of regulatory T cell-mediated CD8 tolerance. Nat. Immunol. 5, 508–515 (2004).

