## Supplementary Information for "Comb-structured mRNA vaccine tethered with short double-stranded RNA adjuvants maximizes cellular immunity for cancer treatment"

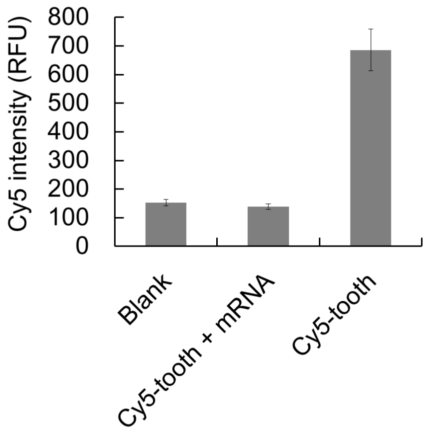


**Figure S1. Hybridization of the immunostimulatory tooth to mRNA.** Comb-structured mRNA with one tooth was prepared from Cy5-labeled 24 nt 5'ppp-RNA (GGUGUGUGUGUGUGUGUGUGUGUG), 43 nt cRNA (with gap sequence of 2 nt) (GGUUCAGGAUGUCCCGCUUCACACACACACACACACACACACC), and *OVA* mRNA (Cy5-tooth + mRNA). After its ultrafiltration with a filter of MWCO 100K, the fluorescence intensity of the flow-through was measured. Cy5-tooth from Cy5-labeled 24 nt 5'ppp-RNA and 43 nt cRNA was used as a control (Cy5-tooth). $n = 4$.

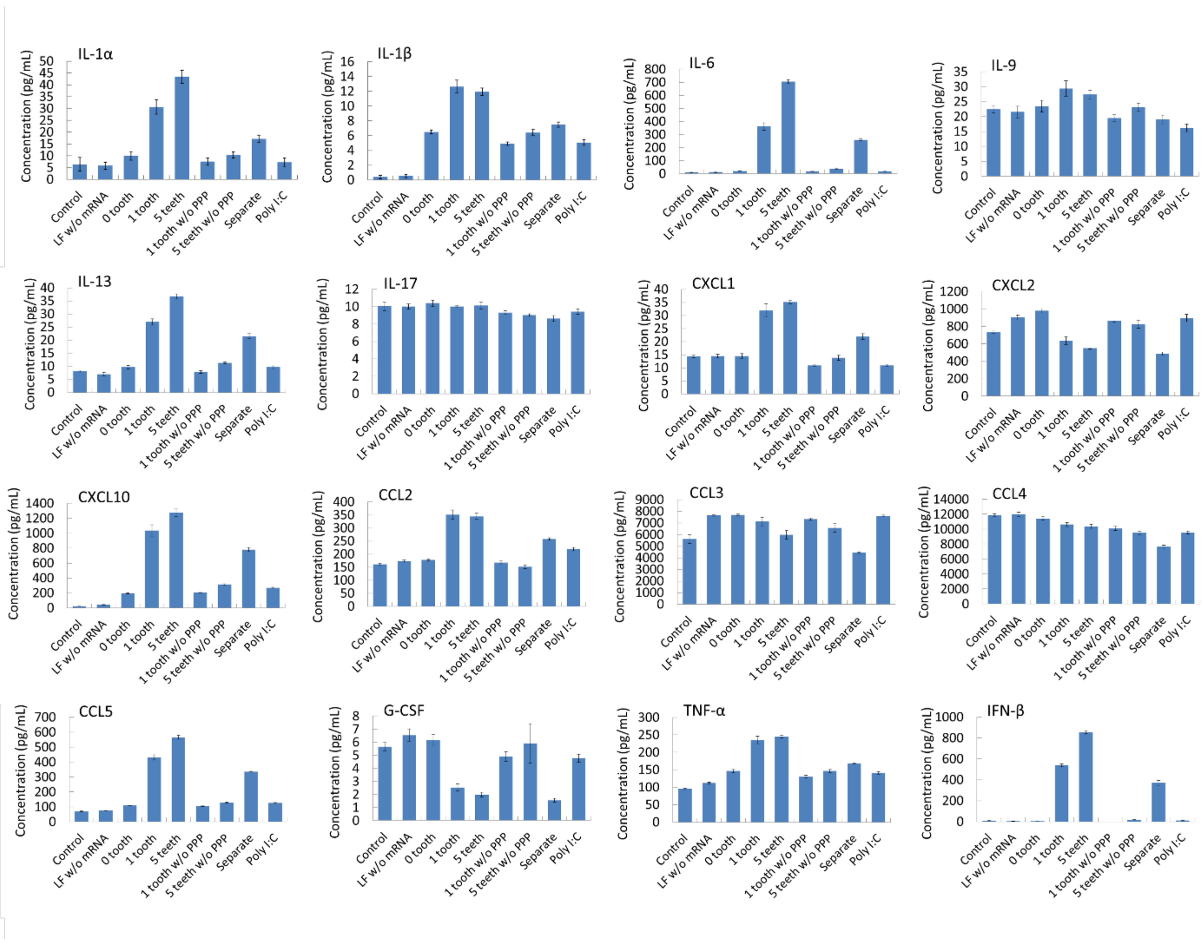


**Figure S2. Immunological profiling of BMDCs after mRNA treatment.** The protein expression profile, shown in **Figure 5**, was presented as the absolute concentration of each protein in the cultured medium. $n = 6$.

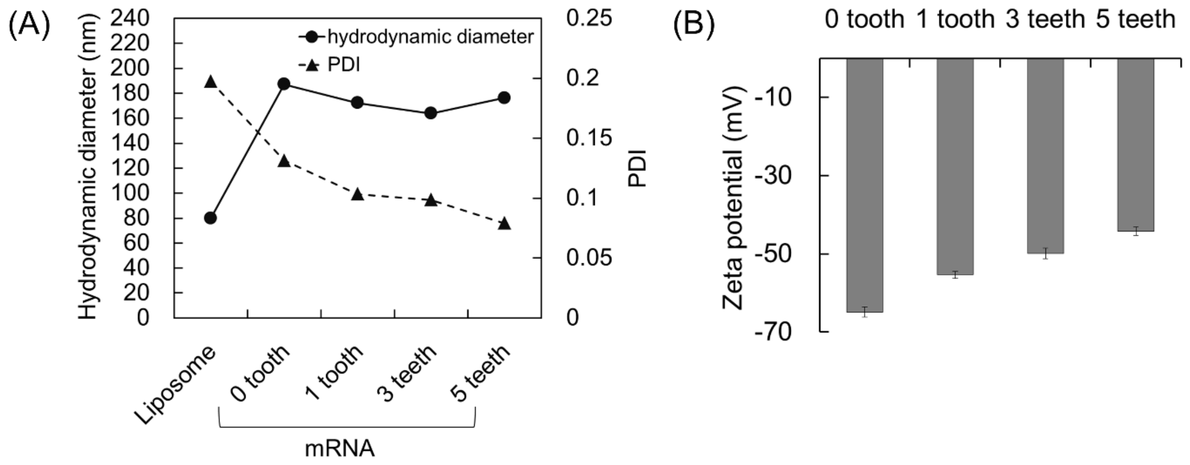


**Figure S3. Physicochemical characterization of lipoplexes.** (**A**) Size and (**B**) ζ-potential of liposome alone and lipoplex loading mRNA with or without immunostimulatory teeth. *n* = 4.

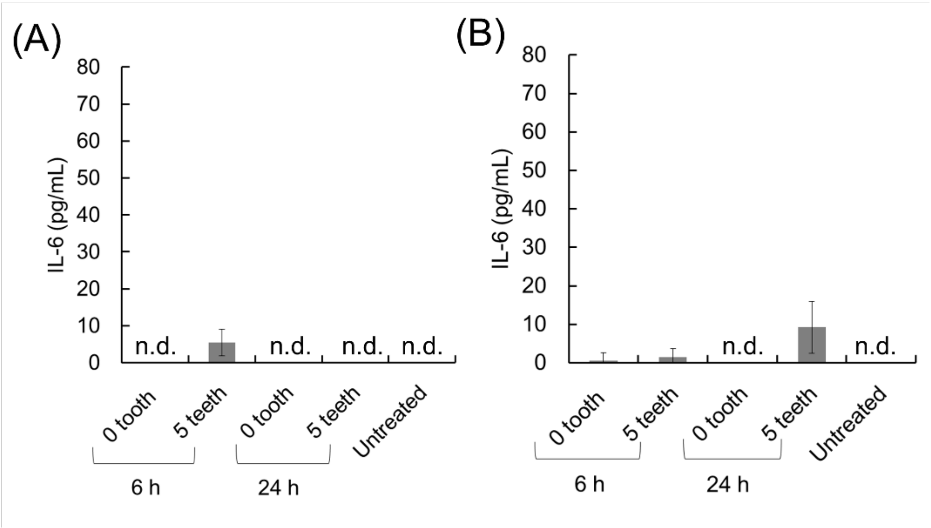


**Figure S4. Safety analyses of PMs in mice.** Serum levels of IL-6 were measured using ELISA 6 h and 24 h after *i.m.* (**A**) *and i.d.* (**B**) injection of PMs. $n = 4$. Note that INF-β was undetected in mouse serum at both time points.

**Supplementary Table S1 Sequences of 5'ppp-RNA and DNA template for 5'ppp-RNA preparation.**

i) 5'ppp-RNA

| | Sequences |
|---|---|
| 24 mer GU-repeat | GGUGUGUGUGUGUGUGUGUGUGUG |
| 40 mer GU-repeat | GGUGUGUGUGUGUGUGUGUGUGUGUGUGUGUGUGUGUGUG |
| 24 mer random | GGUGUGCCUUCUUGGCGUGCGGUG |
| 40 mer random | GGUCCUGGCUUGCCUGUUCCUUCUUCCGUCCGCUCCGUCG |

(ii) DNA template

| | Sequences |
|---|---|
| 24 mer GU-repeat | TAATACGACTCACTATAGGTGTGTGTGTGTGTGTGTGTGTG |
| 40 mer GU-repeat | TAATACGACTCACTATAGGTGTGTGTGTGTGTGTGTGTGTGTGTGTGT GTGTGTGTG |
| 24 mer random | TAATACGACTCACTATAGGTGTGCCTTCTTGGCGTGCGGTG |
| 40 mer random | TAATACGACTCACTATAGGTCCTGGCTTGCCTGTTCCTTCTTCCGTCC GCTCCGTCG |

**Supplementary Table S2 Sequence of cRNA.**

(i) *gLuc* cRNA with 2 nt gap sequence

| Location* | Sequence |
|---|---|
| 60 | CAAACAGAACUUUGACUAACACACACACACACACACACACACC |
| 213 | CUUUGAGCACCUCCAGCAACACACACACACACACACACACACC |
| 244 | CAGCCAGCUUUCCGGGCUACACACACACACACACACACACACC |
| 345 | ACUCUUUGUCGCCUUCGAUCACACACACACACACACACACACC |
| 525 | GCGGCAGCCACUUCUUGUACACACACACACACACACACACACC |

* Locations of the first bases in mRNA hybridized with cRNA are presented as nucleotide numbers from the 5' end. The cRNAs used hybridize to a location 60 in mRNA for 1-tooth mRNA, locations 60, 244, and 525 for 3-teeth mRNA, and locations 60, 213, 244, 345, and 525 for 5-teeth mRNA.

(ii) *gLuc* cRNA with 10 nt gap sequence

| Location* | Sequence |
|---|---|
| 244 | CAGCCAGCUUUCCGGGCUAAAAAAAAACACACACACACACACA CACACC |

* Locations of the first bases in mRNA hybridized with cRNA are presented as nucleotide numbers from the 5' end.

(iii) *fLuc* cRNA

| Location* | Sequence |
|---|---|
| 316 | UCGUUGUAGAUGUCGUUUCACACACACACACACACACACACC |
| 401 | CACGUUCAGGAUCUUCUUUCACACACACACACACACACACACC |
| 694 | AUGGCGGUGUCGGGGAUUUCACACACACACACACACACACACC |
| 882 | GGGUGCUCUUGGCGAAGUUCACACACACACACACACACACACC |
| 1057 | UUGUCGUCGCCCUCGGGUUCACACACACACACACACACACACC |

* Locations of the first bases in mRNA hybridized with cRNA are presented as nucleotide numbers from the first base of the start codon. The cRNAs used hybridize to a location 316 for 1-tooth mRNA and locations 316, 401, 694, 882, and 1057 for 5-teeth mRNA.

(iv) *OVA* cRNA

| Location* | Sequence |
|---|---|
| 252 | GGUUCAGGAUGUCCCGCUUCACACACACACACACACACACACC |
| 332 | GGGCAGGAUGGGGUACCUUCACACACACACACACACACACACC |
| 563 | GUCCUCGUCCUUGAAGGAUCACACACACACACACACACACACC |
| 777 | GCUUCUCGAAGUUGAUGUACACACACACACACACACACACACC |
| 1103 | GGCGAUGUGCUUGAUGCUUCACACACACACACACACACACACC |

* Locations of the first bases in mRNA hybridized with cRNA are presented as nucleotide numbers from the first base of the start codon. The cRNAs used hybridize to a location 252 for 1-tooth mRNA, locations 252, 563, and 1103 for 3-teeth mRNA, and locations 252, 332, 563, 777, and 1103 for 5-teeth mRNA.

(v) *gp100* cRNA

| Location* | Sequence |
|---|---|
| 223 | GUCGUUGAUCACUCUCAUUCACACACACACACACACACACACC |
| 336 | UUGAUGAUUGUGUUGUUUUCACACACACACACACACACACACC |
| 462 | CACACGUACACGAAGCUUUCACACACACACACACACACACACC |
| 598 | CACGUAAGACUGGGAUCUUCACACACACACACACACACACACC |
| 703 | CAGGAAGUGCUUGGUCUUUCACACACACACACACACACACACC |

* Locations of the first bases in mRNA hybridized with cRNA are presented as nucleotide numbers from the 5' end. The cRNAs used hybridize to a location 223 for 1-tooth mRNA and locations 223, 336, 462, 598, and 703 for 5-teeth mRNA.

**Supplementary Table S3 Dynamic light scattering of iLNPs.**

| Tooth number | 0 | 1 |
|---|---|---|
| Size (d, nm) | 88.4 ± 0.3 | 86.3 ± 0.4 |
| Polydispersity index (PDI) | 0.18 ± 0.01 | 0.20 ± 0.01 |

**Supplementary Table S4 Dynamic light scattering of PMs.**

| Tooth number | 0 | 1 | 5 |
|---|---|---|---|
| Size (d, nm) | 78.5 ± 0.8 | 74.5 ± 0.2 | 76.4 ± 0.6 |
| Polydispersity index (PDI) | 0.23 ± 0.01 | 0.21 ± 0.00 | 0.24 ± 0.01 |
